# Branched C7-Substituted 7-Deaza-SAH Analogues Occupy the Entire SAM-Binding Pocket of Mpox Virus VP39 and Dengue Virus NS5 Methyltransferases

**DOI:** 10.64898/2026.07.28.741162

**Authors:** Milan Štefek, Martin Klima, Tomáš Otava, Dominika Chalupska, Milan Dejmek, Radim Nencka, Evzen Boura

**Author notes:** These authors contributed equally (Milan Štefek, Martin Klima).

## Abstract

Viral RNA-cap MTases are attractive targets for antiviral drug development. We previously identified C7-substituted 7-deaza-SAH analogues as potent inhibitors of the mpox virus 2′-O-MTase VP39. Here, we used structure-guided design to develop branched C7-substituted analogues intended to engage multiple hydrophobic regions of the VP39 SAM-binding pocket. The synthesized compounds were characterized using biochemical and crystallographic approaches. Several analogues effectively inhibited VP39, with the most potent compound displaying an IC_50_ in the tens-of-nanomolar range. Crystal structures of VP39 in complex with STM1187 and STM1189 confirmed that the branched aromatic substituents extend towards hydrophobic regions adjacent to the SAM binding site. The precise ligand conformations were strongly influenced by linker geometry and the branching group’s mode of attachment. STM1078 also inhibited DENV3 NS5 MTase with submicromolar potency, and the complex’s structure revealed a conserved binding mode of the SAH-like core accompanied by conformational adaptability of the branched substituent. These results demonstrate how three-dimensional expansion from the 7-deaza position can generate potent inhibitors capable of binding structurally distinct viral MTases.

## Introduction

The 5′ cap is a key feature of eukaryotic mRNA, determining its identity and stability. In addition, it promotes efficient translation [1, 2]. In human cells, canonical mRNA capping begins with the addition of a guanosine linked to the first transcribed nucleotide through a 5′-5′ triphosphate bridge, generating the pre-cap structure. Subsequent *N7*-methylation produces the cap-0, and 2′-O methylation of the first and, in many transcripts, the second transcribed nucleotide produces cap-1 and cap-2 structures, respectively. These modifications protect mRNA from exonucleolytic degradation, promote pre-mRNA splicing, support nuclear export, and stimulate translation initiation through cap-binding proteins [1, 2]. Cap methylation also contributes to innate immune discrimination of self and non-self RNA, because incompletely capped RNAs can be recognized by antiviral sensors and restriction factors, including RIG-I and IFIT proteins. Thus, the mRNA cap is not only a platform for gene expression but also a molecular signature of cellular RNA [3-5].

Viruses must also protect the 5′ end of their mRNA, which in some viruses is also the genomic RNA. Some viruses have evolved alternative strategies; for instance, picornaviruses use the small protein 3B, also known as VPg (viral protein, genome-linked), which is covalently attached to the 5′ end of their genomic RNA. This RNA also serves as mRNA and is translated by an IRES-driven mechanism, bypassing the need for an RNA cap [6]. However, conventional capping, or cap “snatching”, is a widespread strategy used by many medically important viral families, including flaviviruses, coronaviruses, poxviruses, rhabdoviruses, and orthomyxoviruses [4]. Viruses that replicate outside of the nucleus lack access to the host nuclear capping machinery required to produce capped RNA. Therefore, many of them encode their own capping enzymes, including RNA-cap MTases, or use alternative mechanisms, such as VPg-mediated 5′-end protection in the case of the aforementioned picornaviruses. Loss of viral cap methylation can attenuate infection by impairing translation and increasing sensitivity to interferon-stimulated antiviral proteins. These properties make viral RNA-cap MTases targets for antivirals [7-10].

Poxviruses are large double-stranded DNA viruses that replicate predominantly in the cytoplasm and encode much of their own transcription and mRNA-processing machinery [11]. Their mRNAs are capped, methylated, and polyadenylated by virus-encoded enzymes, which allows efficient viral gene expression independently of the host nucleus. The poxviral protein VP39 is a cap-specific 2′-O MTase that converts cap-0 to cap-1 RNA [12]. The 2′ -O MTase activity of VP39 is particularly important for innate immune evasion, since viral RNAs lacking this modification are more efficiently restricted by IFIT-mediated antiviral responses [3, 5].

Viral MTase inhibitor development has expanded substantially in recent years, particularly during the COVID-19 pandemic. The urgent need for antiviral targets during COVID-19 accelerated structural biology, biochemical assay development, and rational inhibitor design against these enzymes [13-18]. These studies showed that viral S-adenosyl-L-methionine (SAM)-dependent MTases can be inhibited by compounds targeting the conserved SAM/S-adenosyl-L-homocysteine (SAH)-binding pocket, by bisubstrate-like inhibitors that also engage the RNA-binding region, or by non-nucleoside molecules exploiting enzyme-specific pockets [9, 13-18].

Importantly, the structural and mechanistic knowledge obtained for coronavirus MTases can be applied to other viral MTases, including monkeypox (mpox) VP39. We previously identified SAH analogues modified at the 7-deaza position as potent inhibitors of mpox VP39 MTase [19, 20]. In the present study, we developed branched bis(aryl)-SAH analogues designed to occupy a larger portion of the VP39 SAM-binding pocket. We identified compounds with submicromolar inhibitory activity and determined crystal structures of VP39 in complex with TO501, STM1187, and STM1189. These structures reveal how linker geometry and branched aromatic substituents control ligand conformation and enable engagement of distinct hydrophobic regions of the SAM/SAH pocket.

## Materials and Methods

### Protein expression and purification

The mpox virus (USA-May22 strain) VP39 MTase and the dengue virus serotype 3 (DENV3) NS5 MTase domain were produced and purified according to previously established protocols [19, 21]. Briefly, both proteins were expressed as fusion constructs with an *N*-terminal 8xHis tag for purification followed by a SUMO tag to enhance solubility and folding. Recombinant expression was carried out in the *E. coli* BL21 DE3 bacterial strain using the LB medium. At an optical density at 600 nm (OD_600_) of 0.6, the production of the target protein was induced by the addition of 250 µM isopropyl-β-D-thiogalactopyranoside (IPTG) and the temperature was lowered to 18°C for 15 hours. Cells were collected by centrifugation, resuspended in the lysis buffer (50 mM Tris.HCl pH 8.0, 300 mM NaCl, 20 mM imidazole, 3 mM β-mercaptoethanol), and disrupted using the Emulsiflex C3 instrument (Avestin). The resulting lysate was precleared by centrifugation at 30,000 g for 30 min and incubated with the HisPur Ni-NTA Superflow agarose (Thermo Fisher Scientific) for 60 min. Then, the resin was extensively washed with the lysis buffer, and the target protein was eluted using the lysis buffer supplemented with 300 mM imidazole. The eluted fusion protein was then treated with Ulp1 protease to remove the 8xHis-SUMO tag. The target proteins were further purified by the size-exclusion chromatography on the HiLoad 16/600 Superdex 75 prep grade column (Cytiva) pre-equilibrated with the size-exclusion buffer composed of 10 mM Tris pH 8.0, 100 mM NaCl, 100 mM LiCl, and 1 mM TCEP in case of mpox virus VP39 protein, or 25 mM Hepes pH 7.5, 500 mM NaCl, 5 % glycerol, and 1 mM TCEP in case of DENV3 NS5 MTase. Fractions containing the purified target proteins were concentrated to 13 mg/mL and directly used for further experiments.

### Crystallization and crystallographic analysis

For structural studies, the VP39 protein or NS5 MTase domain. each at a concentration of 13 mg/ml, was supplemented with a twofold molar excess of the respective ligand (TO501, STM1187, STM1189, or STM1078) and incubated on ice for 1 hour. Protein crystals grew within 3 days at 18 °C in sitting drops consisting of 400 nl of the protein and 400 nl of the reservoir solution using the vapor diffusion method at 291 K. The drops were set up using the Mosquito robot (SPT Labtech). Low-quality crystals generated in the initial crystallization trials were used for preparation of crystallization seeds for further optimization apphroaches. Upon seeding, the final crystals of sufficient quality were prepared using the reservoir solution composed of 200 mM lithium citrate and 15% (w/v) PEG 3.350 for mpox virus VP39 protein, or 200 mM ammonium sulfate, 100 mM MES pH 6.5, and 30% (w/v) PEG 5.000 MME for DENV3 NS5 MTase. Prior to freezing, the crystals were cryo-protected using the reservoir solution supplemented with 20% (v/v) glycerol. Then, the crystals were harvested and flash-cooled in liquid nitrogen. X-ray diffraction data were collected from single crystals on the BL14.1 beamline at the BESSY II electron storage ring operated by the Helmholtz-Zentrum Berlin [22] and on the P13 beamline operated by EMBL Hamburg at the PETRA III electron storage ring (DESY, Hamburg, Germany) [23].

Data integration and scaling were performed using XDS [24]. The structures of the VP39 protein and the NS5 MTase domain in complex with the respective ligands were solved by molecular replacement, using the structures of the VP39/sinefungin complex (pdb entry 8B07) and the NS5 MTase/AT-9010/SAH complex (pdb entry 8BCR), respectively, as search models. The initial model was generated with Phaser v2.8.3 [25] and subsequently improved through automated model refinement with the phenix.refine tool [26] from the Phenix package v1.20.1-4487 [27] and manual model building with Coot v0.9.8.7 [28]. Geometrical restraints for the ligands were prepared using Grade2 v1.8.0 (Global Phasing Ltd.). Statistics for data collection and processing, structure solution and refinement are provided in **SI Table 1**. Structural figures were prepared using the PyMOL Molecular Graphics System v2.5.4 (Schrödinger, LLC). The atomic coordinates and structure factors have been deposited in the Protein Data Bank (https://www.rcsb.org) under the accession codes 30VC, 30VD, 30VE, and 31TF.

### Preparation of RNA substrate for the mpox VP39 MTase assay

A 35-nucleotide RNA transcript carrying an N7-methylguanosine cap structure (m^7^GpppG) was used as the substrate in the 2′-O-methyltransferase assay. The RNA sequence was: m^7^GpppGGGGGAAGCGGGCAUGCGGCCAGCCAUAGCCGAUCA

The transcript was generated by in vitro transcription from the following double-stranded DNA template: 5′-CAGTAATACGACTCACTATAGgggaagcgggcatgcggccagccatagccgatca-3′

Transcription reactions were carried out using the TranscriptAid T7 High-Yield Transcription Kit (Thermo Scientific). Each 500-μl reaction contained 1× TranscriptAid reaction buffer, 8 mM NTPs, 10 μg DNA template, 1× enzyme mix, and 3.5 mM m^7^GpppG cap analog (Jena Bioscience). Following incubation at 37°C for 8 hours, residual template DNA was removed by DNase I (Sigma Aldrich) digestion. The resulting RNA was purified by phenol–chloroform extraction and stored at −20°C until further use. The RNA substrate for the Dengue NS5 MTase was prepared analogously accordingly to our published protocols [29].

### Mpox VP39 2′-O-MTase assay

The assay was performed as previously [19]. Briefly, the MTase reactions were conducted in a final volume of 4 μl using assay buffer composed of 5 mM Tris-HCl (pH 8.0), 1 mM TCEP, 0.1 mg/ml BSA, 0.005% Triton X-100, and 1 mM MgCl_2_. Reaction mixtures contained 200 nM VP39 enzyme, DB RNase inhibitor (DIANA Biotechnologies), and tested compounds at concentrations ranging from 0 to 12.5 μM or 0 to 50 μM for determination of IC_50_ values.

Reactions were initiated by addition of 4 μM SAM together with 4 μM m^7^GpppG-capped RNA substrate. After incubation at 24°C for 45 min, reactions were quenched by adding formic acid to a final concentration of 5%, resulting in a total reaction volume of 6 μl. Samples were analyzed using an Echo acoustic ejection mass spectrometry platform coupled to a Sciex 6500 triple-quadrupole mass spectrometer equipped with an electrospray ionization source. Enzymatic activity was quantified by measuring formation of SAH, the methyl-transfer reaction product. Mass spectrometric detection was performed in multiple-reaction-monitoring (MRM) mode with the ion source temperature maintained at 350°C. Instrument settings included a declustering potential of 20 V, an entrance potential of 10 V, and a collision energy of 28 eV. For analysis, 10 nL of sample was injected into a mobile phase consisting of 70% acetonitrile and 0.1% formic acid flowing at 0.46 mL/min. Quantification of SAH was based on monitoring the characteristic MRM transition m/z 385.1 → 134.1. The dengue MTase assay was performed analogously accordingly to our published protocols [29].

## Results

### Structure guided inhibitor design

In our previous study, we identified several SAH-derived compounds that inhibited the VP39 MTase significantly better than sinefungin (**Fig. 1**) [19]. Most of these compounds contained a large aromatic or heterocyclic substituent connected to the 7-deaza position of the SAH scaffold via an unsaturated alkynyl spacer (such as TO427, **Fig. 2**). The triple-bond-containing linker restricts substituent rotation only to the bond axis. In this study, we prepared another series of SAH-derived compounds containing a large aromatic or heterocyclic substituent connected to the 7-deaza position of the SAH scaffold through fully saturated single-bonded linker and identified four such compounds that inhibited the VP39 MTase significantly better than sinefungin (**Fig. 1**). Analogously to our previous study [14], the benzyl derivative **TO411** was prepared from protected 7-deaza-7-iodo SAH analogue **1** using Negishi cross-coupling reaction. The remaining three SAH derivatives (**TO501, TO512** and **TO739**), which contain an ethylene bridge, were obtained through a two-step synthesis consisting of a Sonogashira cross-coupling reaction with the corresponding (het)arylacetylenes followed by catalytic reduction of the installed triple bond (**SI Fig. 1**).

**Figure 1.**
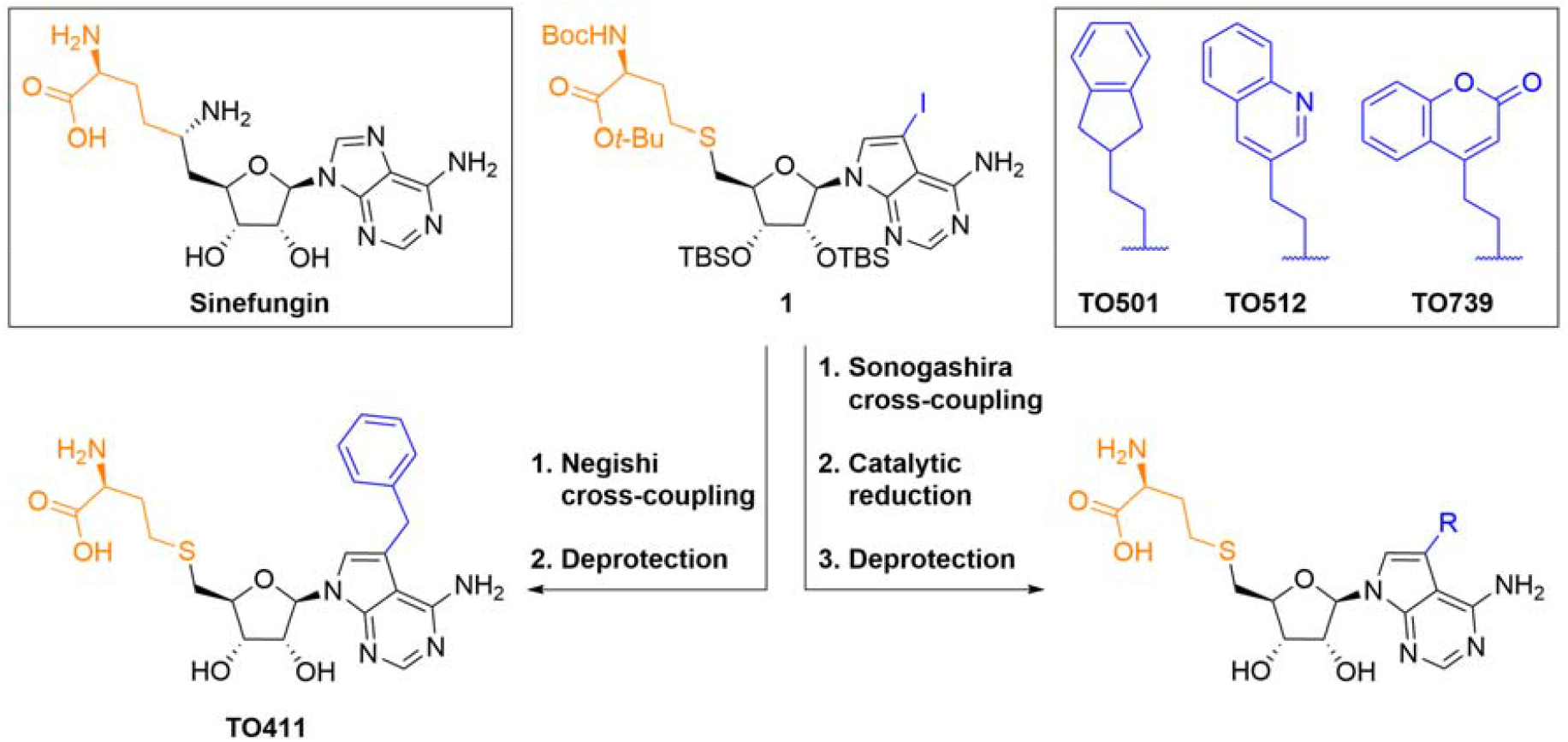
Synthesis of SAH-derived small molecule inhibitors of VP39 containing C7-deaza-substituents linked with a fully saturated single-bonded linker. Experimental details are provided in the Supporting Information.

**Figure 2.**
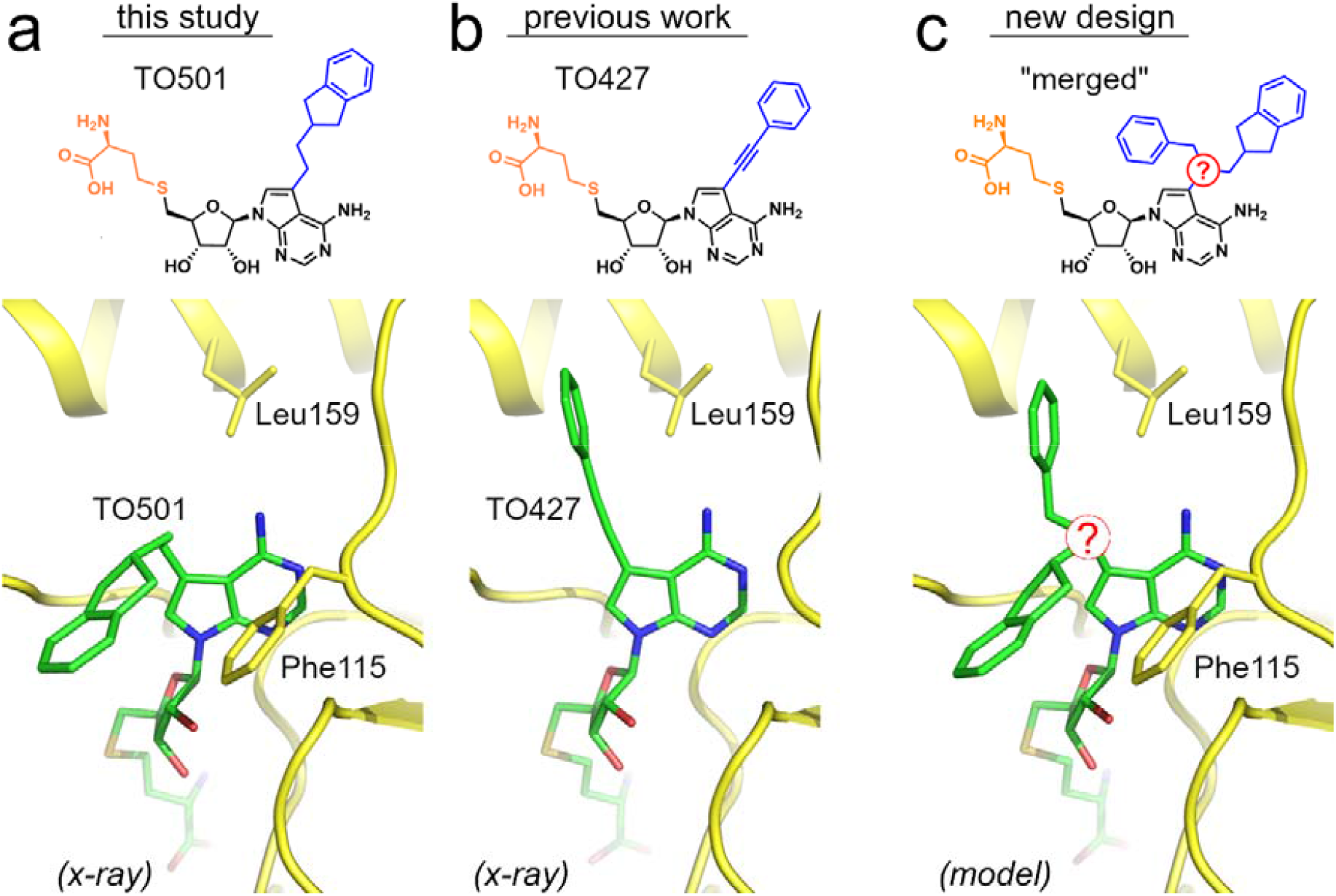
Design of SAH-derived VP39 inhibitors with branched C7-deaza-substituents.

Next, we performed a crystallographic analysis to elucidate the atomic-level details of the interactions between the VP39 protein and the newly identified inhibitors. We succeeded in preparation of diffraction-quality crystals of the VP39 protein in complex with only one of these inhibitors, **TO501**. The crystals belonged to the orthorhombic space group P2_1_2_1_2_1_ and diffracted to 2.3 Å resolution. The respective structure was subsequently solved by molecular replacement and further refined to good R factors and geometry as provided in **SI Table 1**. As anticipated, the **TO501** ligand occupied the SAM/SAH-binding pocket of VP39 (**Fig. 2a**). Interestingly, the hydrophobic 7-deaza-substituent linked via a fully saturated single-bonded linker adopted a bent conformation pointing towards Phe115, significantly different from the conformation of previously reported 7-deaza substituents linked via more rigid triple-bond-containing linkers pointing towards Leu159, such as in the case of **TO427** (**Fig. 2b**). This observation led us to an idea to synthesize another series of SAH-derived compounds containing branched 7-deaza substituents with two large aromatic or heterocyclic substituents that could benefit from hydrophobic or π-π interactions with both Leu159 and Phe115 (**Fig. 2c**).

### Synthesis of branched bis(aryl)-aryl-7-deaza SAH analogues

To evaluate the effect of different *C*7 substituent architectures, five structural classes of SAH analogues were prepared. In all cases, a nitrogen atom was selected as the branching point, while its mode of attachment to the nucleobase was varied. Starting from protected 7-iodo-7-deazaadenosine intermediates **2** and **3** [30, 31], the desired analogues were prepared using synthetic routes tailored to the targeted mode of substituent attachment. (Please, see the Supporting Information for synthetic details). A key step in each route was the introduction of the *C*7 substituent. Three compounds contained a one-atom linker between the branching nitrogen and the nucleobase. The methylene-linked amino derivative **STM1078** was obtained via installation of a carbonyl group followed by reductive amination, the sulfonamide **STM1187** was prepared through a chlorosulfonyl intermediate [32], and the carboxamide **STM1189** was synthesized from a carboxylic acid generated by carbonylative cross-coupling. The SAH analogues **STM1199** and **STM1200**, in which the branching nitrogen is directly attached to the nucleobase, were prepared via Cu-catalyzed C–N bond formation followed by alkylation (**Fig. 3**).

**Figure 3.**
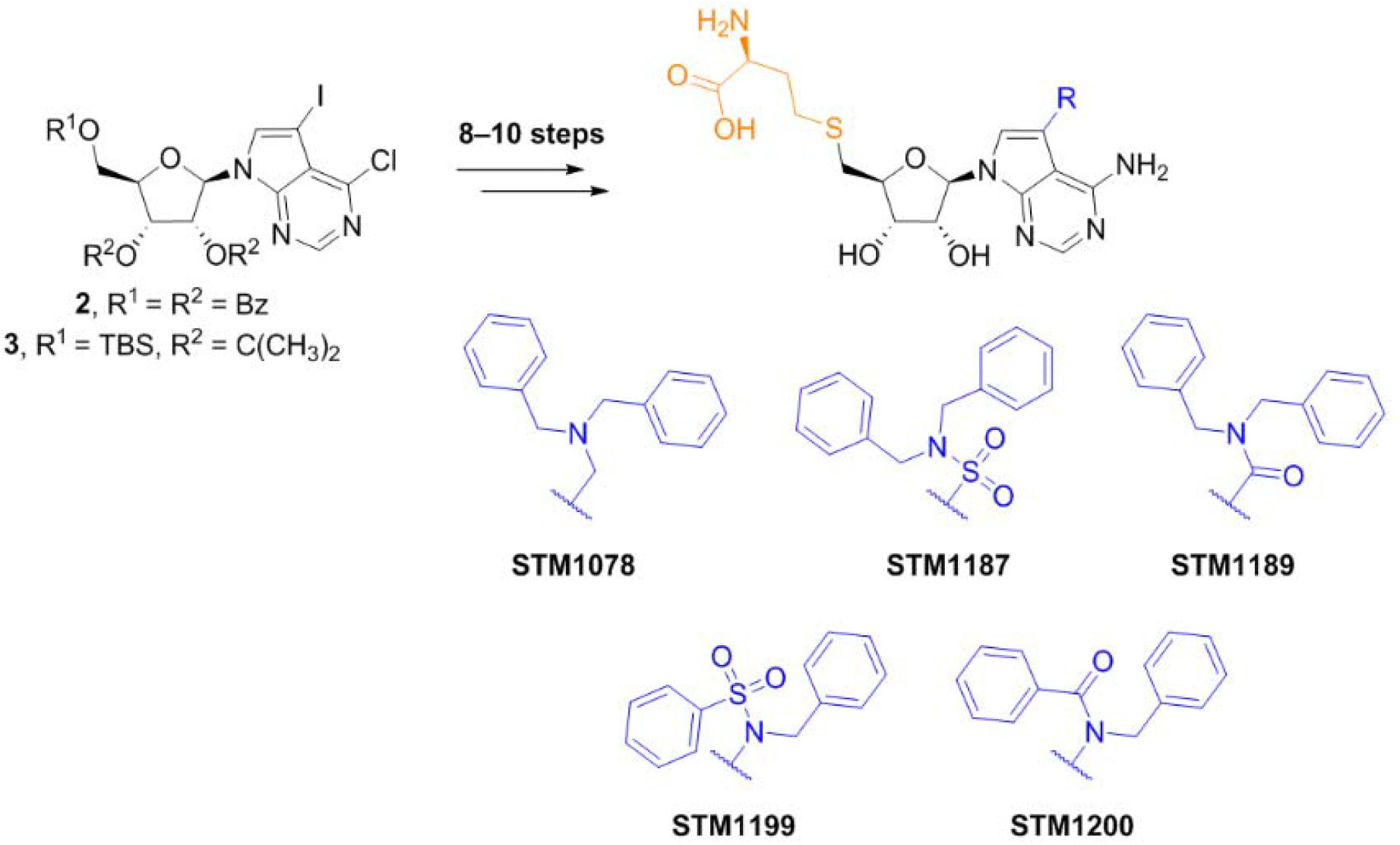
Synthesis of representative C7-substituted SAH analogues featuring different modes of substituent attachment. Experimental details are provided in the Supporting Information.

### Biochemical characterization of the prepared inhibitors

The enzymatic activity of VP39 involves the conversion of Cap-0 RNA to Cap-1 RNA through the transfer of a methyl group from SAM to the 2′-O position of the first transcribed nucleotide. Inhibition of the VP39 2′-O-MTase was assessed by measuring the formation of SAH, the by-product generated during the methylation reaction. Consistent with our previous findings, m7GpppG-capped RNA, which is a preferred substrate of VP39 compared with m7GpppA-capped RNA, was used in all assays [19].

The reference inhibitor sinefungin exhibited an IC_50_value of 2.16 μM [19], whereas compounds **TO411, TO512, TO501**, and **TO739** showed substantially stronger inhibition, with IC_50_values below 0.3 μM.

Testing of the prepared SAH-derived analogues revealed clear differences in VP39 inhibition depending on the mode of *C*7 substituent attachment (**Fig. 4**). The most potent compound was the methylene-linked amine **STM1078**, with an IC_50_ value of 0.06 µM. The sulfonamide-linked analogue **STM1187** also showed strong inhibition (IC_50_ = 0.15 µM), whereas the carboxamide **STM1189** displayed lower, but still submicromolar, potency (IC_50_ = 0.62 µM).

**Figure 4.**
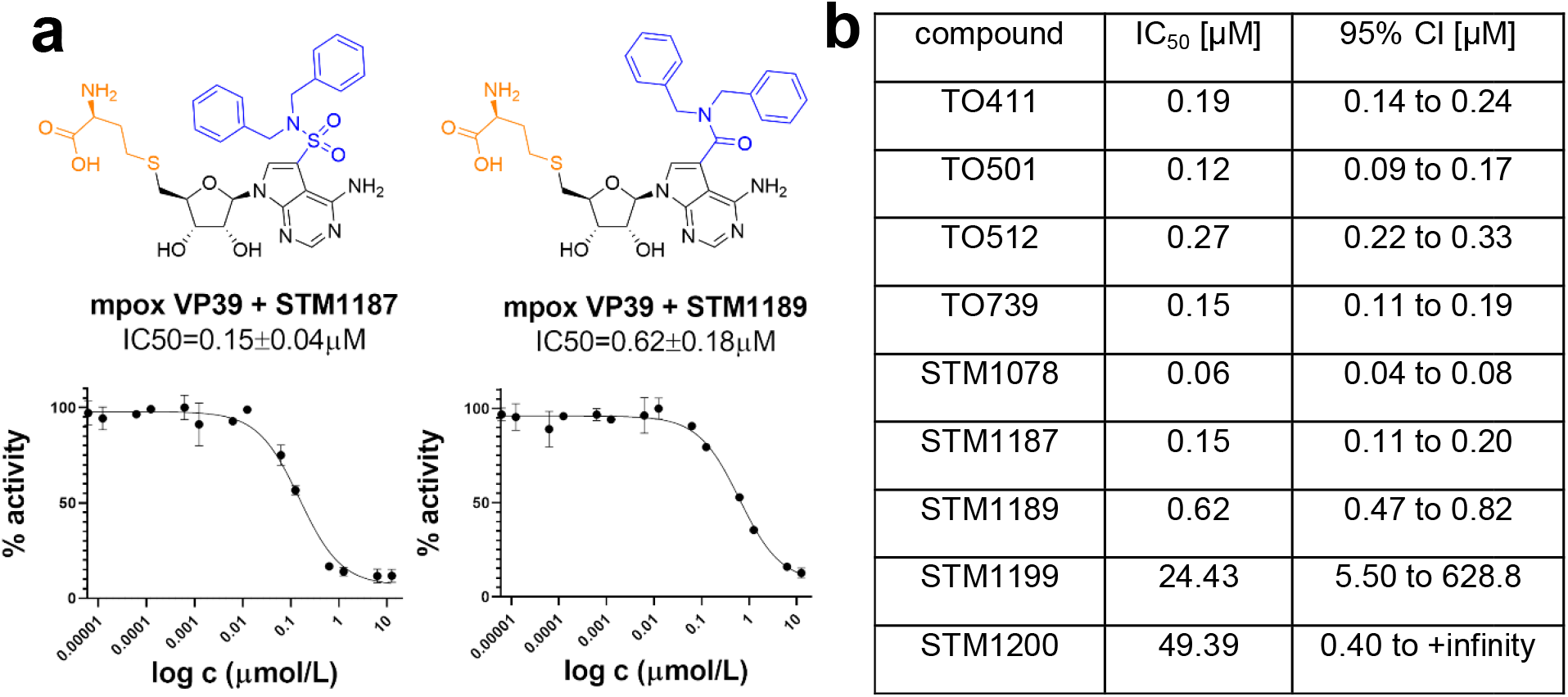
Inhibitory activity of SAH-derived compounds against mpox virus VP39 2′-O-MTase. **a**) The inhibitory activity of the indicated compounds was evaluated using an in vitro VP39 methyltransferase assay; data for **ST 1187** and **STM1189** are shown. IC_50_ values are reported together with their corresponding 95% confidence intervals (95% CI) and were determined by nonlinear regression analysis using GraphPad Prism 8.0.1. IC_50_ values and inhibition curves fitted to data from the MTase inhibition assay are shown. Data points and IC50 values are presented as mean values ± SEM (n = 2 independent measurements). **b**) IC_50_ values of the other compounds prepared in this study.

In contrast, analogues in which the branching nitrogen was directly attached to the nucleobase, **STM1199** and **STM1200**, showed only weak inhibition.

### Crystallographic analysis of the bis(aryl)-SAH analogues bound to VP39

Finally, we carried out a structure analysis of the VP39 protein in complex with these novel compounds containing branched 7-deaza substituents to verify the theoretical model of their mode of binding. We succeeded in preparation of diffraction-quality crystals for two compounds, STM1187 and STM1189. In both cases, the crystals belonged to the monoclinic space group P2_1_ and diffracted to 1.6 Å and 2.6 Å resolution, respectively (**SI Table 1**). The structures revealed that the aromatic moieties of the branched 7-deaza substituents of the compounds pointed towards Phe115 and Leu159 as anticipated; however, their precise conformations were strongly dependent on how the branching of the substituents was chemically achieved **(Fig. 5)**.

**Figure 5.**
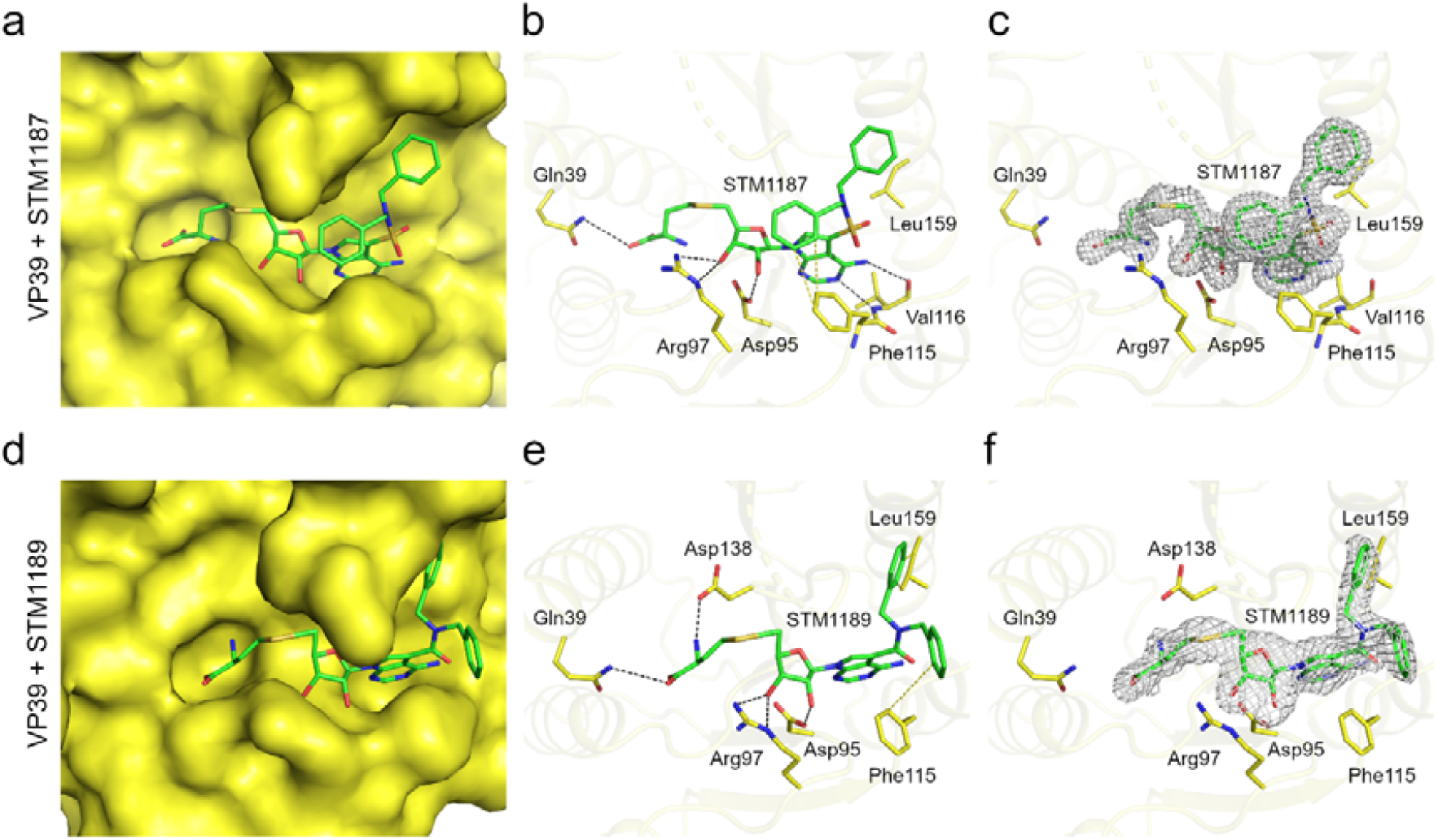
Crystal structure of mpox virus VP39 in complex with the STM1187 and STM1189 compounds. **a-c**, Structural details of the VP39-STM1187 interaction. The ligand is shown in stick representation and colored according to elements: carbon, green; nitrogen, blue; oxygen, red; sulphur, orange. In (**a**), the VP39 protein is shown in surface representation and colored yellow. In (**b**), selected hydrogen bonds and hydrophobic interactions involved in ligand binding are indicated as dashed black and yellow lines, respectively. In (**c**), the Fo-Fc omit map contoured at 2Å is shown around the ligand. **d-f**, Structural details of the VP39-STM1189 interaction depicted as in (**a-c**).

#### Inhibition of flaviviral MTases

We next asked whether branched bis(aryl) 7-deaza SAH analogues can inhibit other viral MTases. We selected the flaviviral NS5 MTase from dengue virus serotype 3 (DENV3), given the medical importance of flaviviruses and our established expertise with these enzymes [29, 33]. Testing of the compound series showed that several analogues inhibited DENV3 NS5 MTase. Among them, **STM1078** displayed submicromolar activity, with an IC_50_ value of 0.49 ± 0.28 μM (**Figure 5**).

To examine the binding mode of **STM1078**, we co-crystallized it with the DENV3 NS5 MTase domain. The resulting diffraction-quality crystals belonged to the hexagonal space group P6_5_ and diffracted to 2.4 Å resolution (**SI Table 1**). The asymmetric unit contained eight protein molecules, each with one **STM1078** molecule bound in the conserved SAM/SAH-binding pocket. The SAH-like core adopted the expected conserved binding mode, whereas the branched 7-deaza substituent showed modest conformational variability among the chains. These data show that branched 7-deaza SAH analogues can bind the SAM pockets of structurally distinct viral MTases and may therefore provide a useful starting point for broader-spectrum antiviral MTase inhibitors.

## Discussion

Based on previous crystal structures of mpox virus VP39 in complex with SAH analogues, we designed branched C7-substituted 7-deaza-SAH analogues. We synthesized these compounds and evaluated them biochemically and structurally. Biochemical evaluation identified several potent inhibitors, with STM1078 and STM1187 displaying low-nanomolar to submicromolar activity against the VP39 MTase. During their design, we hypothesized that these compounds would occupy a larger portion of the SAM-binding pocket of VP39. This was confirmed by the crystal structures that we solved. Moreover, the structures revealed how linker geometry and the mode of substituent attachment determine ligand conformation and engagement of distinct hydrophobic regions of the pocket.

SAH analogues can inhibit several MTases, with the pan-MTase inhibitor sinefungin being a prominent example. However, sinefungin is chemically very similar to SAM; it differs by only a few atoms (the sulfonium methyl-donor centre, –S^+^(CH_3_)–, is replaced by a carbon atom bearing an amino group, –CH(NH_2_)–.). In contrast, the STM compounds possess a (pseudo)adenine ring bearing a large substituent. We have previously shown that SAH analogues bearing a large substituent at the 7-deaza position are not toxic to human cells [20], and we have observed that 7-deaza SAH analogues can inhibit multiple viral MTases [34, 35]. This motivatated us to screen the STM compounds against the flaviviral NS5 MTase. Given the large 7-deaza substituent, which was specifically designed for the active site of the mpox VP39, we did not actually expect to observe any activity against the flaviviral MTase. However, STM1078 proved to be a submicromolar inhibitor of the flaviviral MTase (Fig. 6) demonstrating that this scaffold is not restricted to poxviral VP39 but can also, in principle, target other viral MTases.

**Figure 6.**
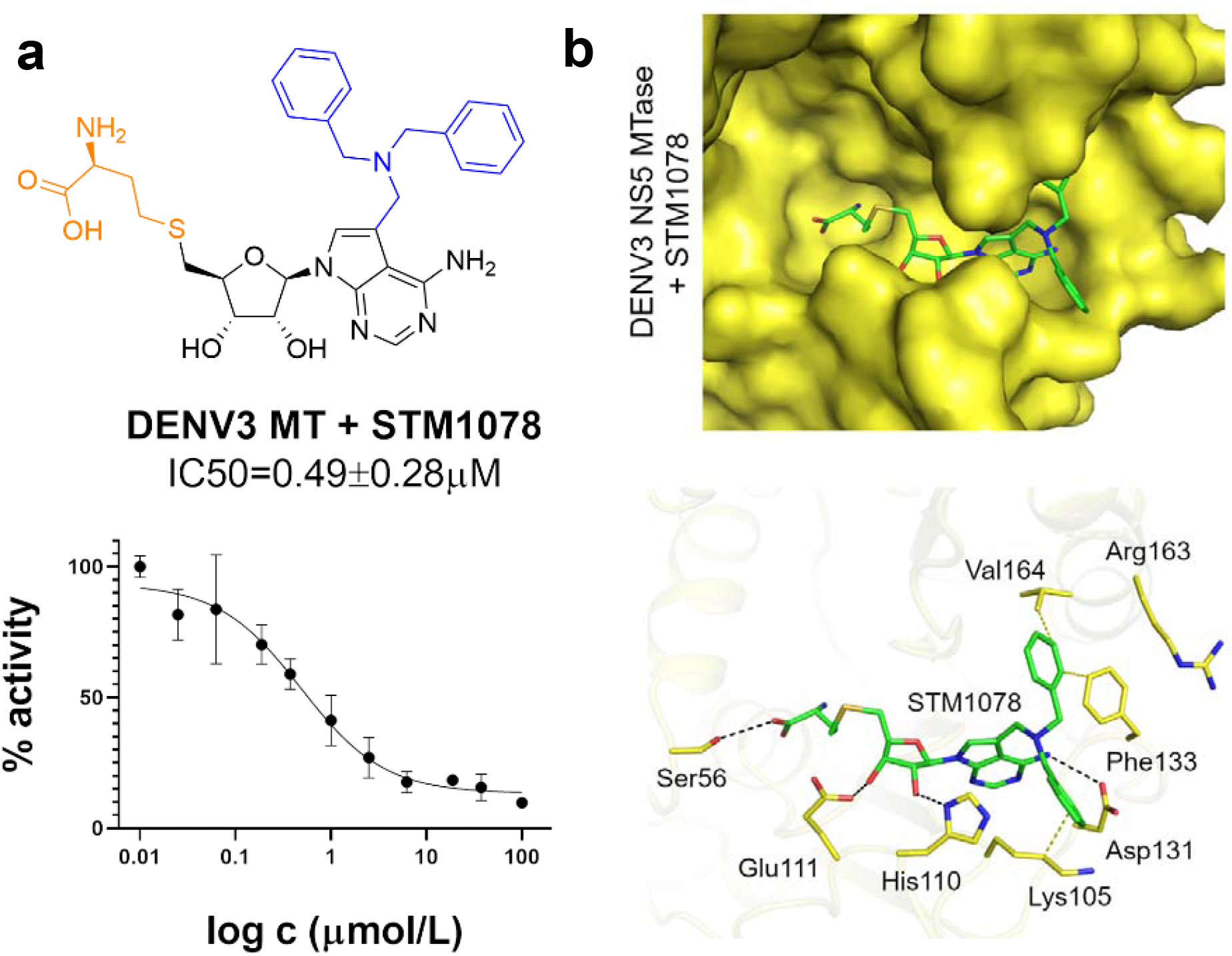
Analysis of STM1078 binding to DENV3 MTase. **a)** IC_50_ determination **b**) Crystallographic analysis.

Indeed, it has been demonstrated that coronavirus MTases can be inhibited by structurally diverse compounds targeting the conserved SAM/SAH-binding pocket. These include SAM- or SAH-derived inhibitors of nsp14 [14, 18], as well as adenine dinucleosides and sulfonamide-based bisubstrate analogues that extend towards the RNA-binding region and engage both the cofactor- and substrate-binding sites [15, 16]. Chemically distinct non-nucleoside inhibitors of nsp14 have also been reported [9]. In contrast to bisubstrate inhibitors, the STM compounds do not extend towards the RNA- or cap-binding region but expand laterally from the 7-deaza nucleobase into hydrophobic regions surrounding the cofactor-binding pocket. These strategies therefore exploit different extensions of the conserved SAM/SAH-binding site and may be combined in future efforts to improve potency and selectivity. A remaining challenge is the translation of potent biochemical inhibition into cellular antiviral activity. The polar SAH-derived scaffold may limit membrane permeability, and the use of prodrugs may therefore be necessary to develop effective antiviral compounds.

## Acknowledgement

We would like to thank EMBL Hamburg and the Helmholtz-Zentrum Berlin für Materialien und Energie for the allocation of synchrotron radiation beamtime. The synchrotron data were collected at beamline P13 operated by EMBL Hamburg at the PETRA III storage ring (DESY, Hamburg, Germany) and at beamline BL14.1 at the BESSY II electron storage ring operated by the Helmholtz-Zentrum Berlin. We would like to thank Kirill Kovalev and Tatjana Barthel for their assistance in using the beamlines. The research was funded by the project “New Technologies for Translational Research in Pharmaceutical Sciences”/NETPHARM, project ID CZ.02.01.01/00/22_008/0004607 (co-funded by the European Union).

## Notes

### Competing Interest Statement

The authors have declared no competing interest.

