## Supplementary data for "Branched C7-Substituted 7-Deaza-SAH Analogues Occupy the Entire SAM-Binding Pocket of Mpox Virus VP39 and Dengue Virus NS5 Methyltransferases"

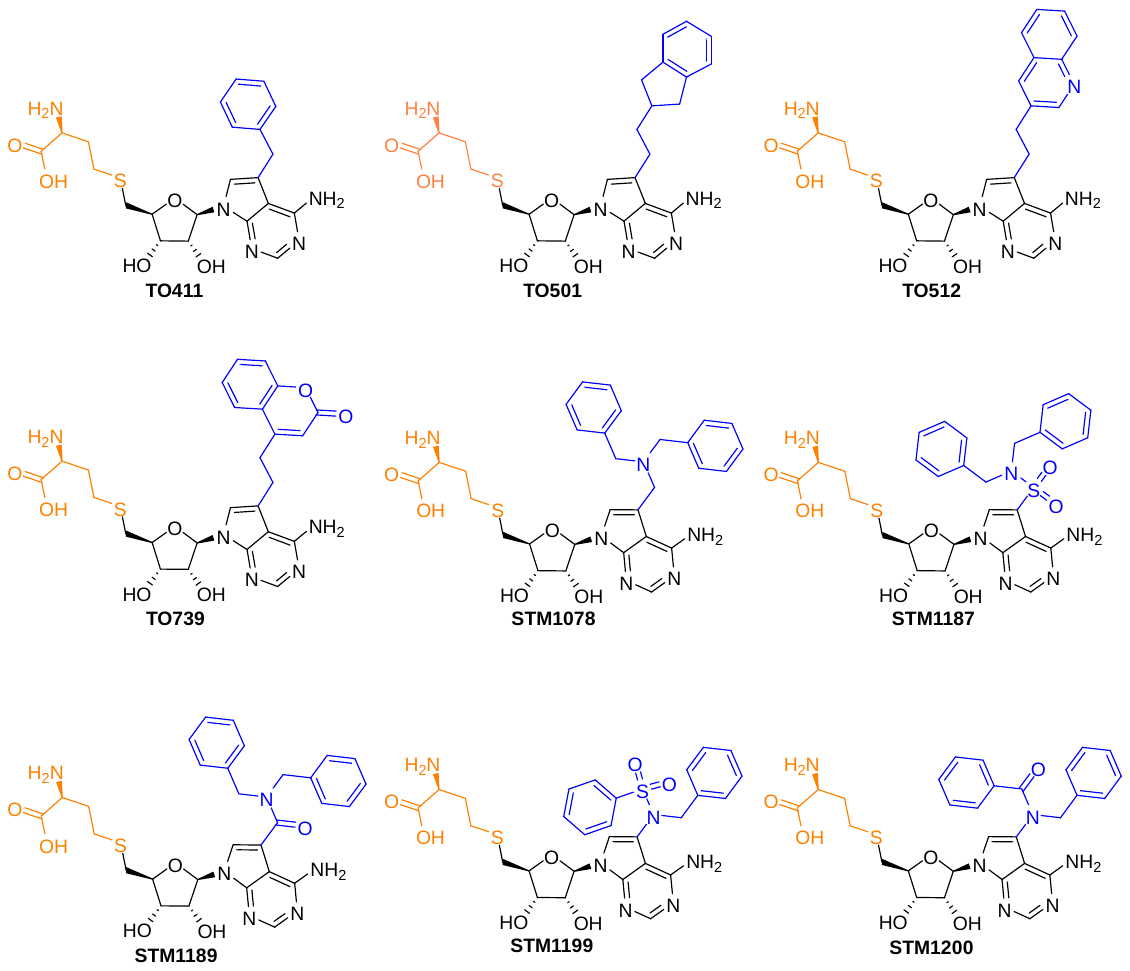

**SI Figure 1** – Expanded chemical structures of the compounds represented by generalized structures in **Figures 1** and **4**.

| **Crystal** | | **VP39 +  TO501** | **VP39 + STM1187** | **VP39 + STM1189** | **DENV3(MT) + STM1087** |
| --- | --- | --- | --- | --- | --- |
| PDB accession code | | 30VC | 30VD | 30VE | 31TF |
| **Data collection and processing** | | | | | |
| Space group | | P 21 21 21 | P 21 | P 21 | P 65 |
| Cell dimensions | a, b, c (Å) | 52.0 84.4 154.9 | 55.9 111.4 56.0 | 62.0 118.5 62.1 | 109.9 109.9 351.6 |
|  | α, β, γ (°) | 90.0 90.0 90.0 | 90.0 98.8 90.0 | 90.0 92.5 90.0 | 90.0 90.0 120.0 |
| Resolution range (Å) | | 49.29 - 2.30 (2.38 - 2.30) | 34.61 - 1.62 (1.68 - 1.62) | 35.88 - 2.64 (2.73 - 2.64) | 47.59 - 2.37 (2.46 - 2.37) |
| No. of unique reflections | | 31,106 (3,044) | 85,830 (8,562) | 46,679 (4,681) | 96,906 (9,674) |
| Completeness (%) | | 99.9 (99.5) | 100.0 (100.0) | 99.8 (99.7) | 99.9 (99.7) |
| Multiplicity | | 19.8 (15.8) | 32.3 (21.9) | 6.8 (7.0) | 20.7 (15.3) |
| Mean I/σ(I) | | 10.92 (1.40) | 15.59 (2.79) | 10.54 (1.98) | 17.99 (2.10) |
| Wilson B factor (Å^2^) | | 32.90 | 20.93 | 29.18 | 48.15 |
| R-merge | | 0.2765 (2.465) | 0.2151 (1.216) | 0.1685 (1.109) | 0.1392 (1.334) |
| R-meas | | 0.2839 (2.548) | 0.2185 (1.247) | 0.1824 (1.197) | 0.1426 (1.38) |
| CC1/2 (%) | | 99.7 (60.6) | 99.7 (63.2) | 99.6 (53.5) | 99.8 (48.3) |
| CC* (%) | | 99.9 (86.9) | 99.9 (88.0) | 99.9 (83.5) | 99.9 (80.7) |
| **Structure solution and refinement** | | | | | |
| R-work (%) | | 21.49 (28.81) | 23.41 (30.96) | 21.52 (26.62) | 23.18 (32.25) |
| R-free (%) | | 24.68 (30.90) | 25.50 (35.68) | 23.85 (28.60) | 26.66 (36.29) |
| CC-work (%) | | 95.4 (76.2) | 91.7 (71.9) | 84.1 (48.7) | 95.1 (60.6) |
| CC-free (%) | | 92.7 (85.4) | 91.6 (69.6) | 83.5 (34.8) | 92.8 (59.6) |
| R.m.s.d. | bonds (Å) | 0.164 | 0.167 | 0.194 | 0.181 |
|  | angles (°) | 1.34 | 1.01 | 2.50 | 3.20 |
| Average B factor (Å^2^) | overall | 42.19 | 26.72 | 34.36 | 60.62 |
|  | protein | 42.45 | 26.45 | 34.37 | 60.47 |
|  | ligands | 38.78 | 27.45 | 41.64 | 70.94 |
|  | solvent | 34.99 | 31.37 | 29.89 | 50.92 |
| Clashscore | | 2.52 | 2.99 | 1.50 | 1.95 |
| Ramachandran (%) | favored | 98.50 | 99.07 | 99.81 | 98.67 |
|  | allowed | 1.50 | 0.93 | 0.19 | 01.33 |
|  | outliers | 0.00 | 0.00 | 0.00 | 0.00 |

**SI Table 1** – Statistics for data collection and processing, structure solution and refinement of the crystal structures of the mpox virus VP39 MTase in complex with the TO501, STM1187, and STM1189 compounds, and Dengue virus-3 NS5 MTase domain in complex with STM1078. Numbers in parentheses refer to the highest resolution shell. R.m.s.d., root-mean-square deviation.

### General information for synthetic part

Synthesis of methyltransferase inhibitors **TO411** and **TO512** was performed according to published procedures.[1]

Starting compounds and reagents were purchased from commercial suppliers (Sigma-Aldrich, Fluorochem, Acros Organics, Carbosynth) and used without further purification. Acetonitrile and 1,4-dioxane were dried using activated 3Å molecular sieves, and DCM was distilled from P_2_O_5_ and kept over 4Å molecular sieves.

Analytical High-Performance Liquid Chromatography (HPLC) and low-resolution mass spectra were measured on a Waters Ultra-high Performance Liquid Chromatography-Mass Spectrometry (UPLC-MS) system consisting of a Waters UPLC H-Class Core System, a UPLC photodiode array (PDA) detector, Waters QDa mass spectrometer, and MassLynx mass spectrometry software. HPLC conditions used were: column Waters CORTECS UPLC C18 column, 1.6 µm, 2.1 × 50 mm, or Waters Acquity UPLC BEH C18 1.7 µm, 2.1 mm × 100 mm; LC method: H_2_O/CH_3_CN, 0.1% formic acid as a modifier, gradient 0–100%). Analytical Thin-Layer Chromatography (TLC) was performed on silica gel-precoated aluminium plates with a fluorescent indicator (Merck 60 F_254_).

For normal flash column chromatography (VWR International Silica gel 60, particle size 0.040–0.063 mm) as well as for reversed-phase flash column chromatography (C18 RediSep Rf columns), a Combiflash® Rf from Teledyne ISCO was used.

^1^H and ^13^C NMR spectra for the reported compounds were recorded on a Bruker Avance III^TM^ HD 400 instrument (400.0 MHz for ^1^H, 101 MHz for ^13^C and 377 MHz for ^19^F) using inverse broadband probe with ATM module (5 mm BBO-1H Z-GRD), Bruker Avance III^TM^ HD 400 instrument with broadband PRODIGY cryoprobe with ATM module (5 mm CPBBO BB-1H/19F/D Z-GRD), or Bruker AVANCE III^TM^ HD 500 MHz spectrometer (500.0 MHz for ^1^H, 125.7 MHz for ^13^C and 470 MHz for ^19^F). Chemical shifts (δ) and coupling constants (*J*) are expressed in ppm and Hz, respectively. The NMR experiments were performed in DMSO-*d*_6_ or CDCl_3_ and referenced to the solvent signal (DMSO-*d*_6_: δ 2.50 for ^1^H NMR and 39.70 for ^13^C NMR; CDCl_3_: δ 7.26 for ^1^H NMR and 77.16 for ^13^C NMR). Shifts of ^1^H and ^13^C, observed only in 2D spectra, are marked with an asterisk (*).

High-resolution mass spectrometry (HRMS) analyses were carried out on an LTQ XL Orbitrap XL (Thermo Fisher Scientific) using electrospray ionization (ESI).

### Synthetic procedures

***tert*-Butyl *S*-(((2*S*,3*R*,4*R*,5*R*)-5-(5-((1*H*-inden-2-yl)ethynyl)-4-amino-7*H*-pyrrolo[2,3-d]pyrimidin-7-yl)-3,4-bis((*tert*-butyldimethylsilyl)oxy)tetrahydrofuran-2-yl)methyl)-*N*-(*tert*-butoxycarbonyl)-l-homocysteinate (4)**

Triethylamine (110 µL, 0.79 mmol, 3 eq) was added to a mixture of protected 7-deaza-7-iodo SAH analogue **1** (236 mg, 0.26 mmol), prepared according to a reported procedure,[1] 2-ethynyl-1*H*-indene (74 mg, 0.53 mmol, 2 eq), CuI (15 mg, 0.08 mmol, 0.3 eq) and Pd(PPh_3_)_4_ (31 mg, 0.03 mmol, 0.1 eq) in dry THF (6 mL) and the suspension was stirred under argon atmosphere at 60 °C for 2 h. After cooling to RT, the mixture was diluted with EtOAc (100 mL) and washed with water (30 mL). Organic layer was dried over anhydrous Na_2_SO_4_, and the solvent was removed in vacuo. The crude mixture was subjected to FCC (3–35% of EtOAc in DCM)
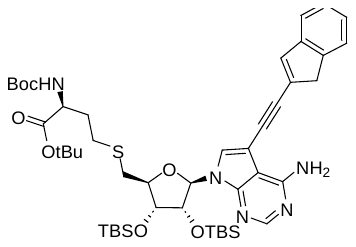
, furnishing derivative **4** (257 mg, 80%) as a brownish foam. **^1^H NMR** (400 MHz, DMSO-*d*_6_) δ 8.16 (s, 1H), 7.91 (s, 1H), 7.49–7.42 (m, 2H), 7.33–7.27 (m, 2H), 7.24 (td, *J* = 7.3, 1.3 Hz, 1H), 7.17 (d, *J* = 7.9 Hz, 1H), 6.14 (d, *J* = 7.1 Hz, 1H), 4.81 (dd, *J* = 7.2, 4.3 Hz, 1H), 4.20 (dd, *J* = 4.2, 1.4 Hz, 1H), 3.98 (t, *J* = 7.1 Hz, 1H), 3.92 (td, *J* = 8.4, 5.7 Hz, 1H), 3.72 (s, 2H), 3.05 (dd, *J* = 13.8, 8.4 Hz, 1H), 2.86 (dd, *J* = 13.8, 6.3 Hz, 1H), 2.65–2.42* (m, 2H), 1.93–1.77 (m, 1H), 1.37 (s, 18H), 0.93 (s, 9H), 0.69 (s, 9H), 0.16 (s, 3H), 0.13 (s, 3H), -0.09 (s, 3H), -0.38 (s, 3H). **^13^C NMR** (101 MHz, DMSO-*d*_6_) δ 171.6, 157.8, 155.7, 153.2, 150.6, 143.9, 142.9, 136.7, 127.1, 127.0, 126.9, 125.9, 123.8, 121.5, 102.2, 95.8, 89.2, 87.7, 86.6, 84.8, 80.5, 78.3, 74.7, 53.5, 42.3, 33.6, 31.2, 28.4, 28.3, 27.8, 26.0, 25.7, 17.9, 17.6, -4.4, -4.5, -4.6, -5.3. **LRMS** (ESI) [M + H]^+^ m/z: 906.5.

***tert*-Butyl *S*-(((2*S*,3*R*,4*R*,5*R*)-5-(4-amino-5-(2-(2,3-dihydro-1*H*-inden-2-yl)ethyl)-7*H*-pyrrolo[2,3-d]pyrimidin-7-yl)-3,4-bis((*tert*-butyldimethylsilyl)oxy)tetrahydrofuran-2-yl)methyl)-*N*-(*tert*-butoxycarbonyl)-l-homoczysteinate (5)**

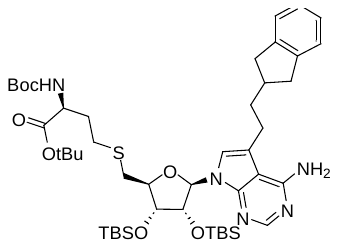
SAH analogue **4** (200 mg, 0.22 mmol) and 10% Pd/C (53 mg, 0.04 mmol, 0.2 eq) were suspended in a 1:1 mixture of EtOH and EtOAc (14 mL). The mixture was stirred vigorously under an H_2_ atmosphere at room temperature for 18 h. After completion, the catalyst was removed by filtration through a pad of Celite, and the filtrate was concentrated under reduced pressure. The crude material was purified by FCC (3–35% of EtOAc in DCM) to give the desired compound **5** (160 mg, 80%) as a white foam. **^1^H NMR** (400 MHz, DMSO-*d*_6_) δ 8.02 (s, 1H), 7.21–7.12 (m, 4H), 7.12–7.05 (m, 2H), 6.53 (s, 2H), 6.09 (d, *J* = 7.3 Hz, 1H), 4.75 (dd, *J* = 7.3, 4.4 Hz, 1H), 4.17 (dd, *J* = 4.4, 1.4 Hz, 1H), 3.97–3.87 (m, 2H), 3.09–2.94 (m, 3H), 2.91–2.73 (m, 3H), 2.65–2.43* (m, 4H), 1.90–1.73 (m, 4H), 1.37 (s, 9H), 1.36 (s, 9H), 0.92 (s, 9H), 0.64 (s, 9H), 0.14 (s, 3H), 0.11 (s, 3H), -0.14 (s, 3H), -0.41 (s, 3H). **^13^C NMR** (101 MHz, DMSO-*d*_6_) δ 171.6, 157.8, 155.7, 151.7, 151.6, 143.2, 126.1, 124.4, 118.9, 115.8, 102.4, 86.0, 84.2, 80.5, 78.3, 74.7, 73.9, 53.5, 38.9, 38.9, 36.1, 33.7, 31.2, 28.4, 28.3, 27.8, 26.0, 25.7, 24.8, 17.9, 17.7, -4.4, -4.4, -4.6, -5.3. **LRMS** (ESI) [M + H]^+^ m/z: 912.5.

***S*-(((2*S*,3*S*,4*R*,5*R*)-5-(4-Amino-5-(2-(2,3-dihydro-1*H*-inden-2-yl)ethyl)-7*H*-pyrrolo[2,3-d]pyrimidin-7-yl)-3,4-dihydroxytetrahydrofuran-2-yl)methyl)-l-homocysteine (TO501)**

To a solution of protected SAH analogue **5** (145 mg, 0.16 mmol) in a mixture of TFA and water (9:1, 4 mL) at 0 °C. After stirring for 3 h at RT, the mixture was concentrated under reduced pressure, and the residue was co-evaporated twice with MeOH. Then, the crude mixture was re-dissolved in MeOH (2 mL) and neutralized with NH_4_OH. The volatiles were removed under reduced pressure, and the residue was purified by RP-FCC (3–40% of MeOH in water), providing the final compound **TO501** (71 mg, 0.14 mmol, 85%) as a white solid.
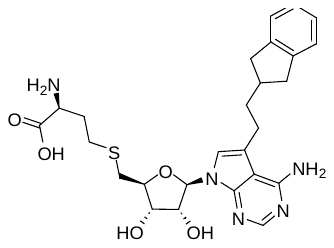
 **^1^H NMR** (500 MHz, DMSO-*d*_6_) δ 8.03 (s, 1H), 7.18 (dt, *J* = 7.1, 3.4 Hz, 2H), 7.12 (s, 1H), 7.11–7.06 (m, 2H), 6.56 (s, 2H), 6.04 (d, *J* = 5.9 Hz, 1H), 5.43 (bs, 2H), 4.41 (t, *J* = 5.7 Hz, 1H), 4.02 (dd, *J* = 5.4, 3.8 Hz, 1H), 3.92 (td, *J* = 6.3, 3.7 Hz, 1H), 3.27 (dd, *J* = 7.2, 5.3 Hz, 1H), 3.09–3.00 (m, 2H), 2.90–2.80 (m, 3H), 2.71 (dd, *J* = 13.6, 6.5 Hz, 1H), 2.65–2.56 (m, 4H), 2.55–2.42* (m, 1H), 2.04–1.94 (m, 1H), 1.86–1.74 (m, 3H). **^13^C NMR** (126 MHz, DMSO-*d*_6_) δ 169.8, 157.9, 151.7, 151.4, 143.4, 126.2, 124.5, 118.6, 115.9, 102.3, 86.5, 82.7, 73.2, 72.8, 53.2, 53.1, 38.9, 38.9, 36.0, 34.3, 31.6, 28.4, 24.9. **HRMS** (ESI) m/z: [M + H]^+^ (C_26_H_33_N_5_O_5_S) calculated 526.213, found 526.213.

***tert*-Butyl *S*-(((2*S*,3*R*,4*R*,5*R*)-5-(4-amino-5-((2-oxo-2*H*-chromen-4-yl)ethynyl)-7*H*-pyrrolo[2,3-d]pyrimidin-7-yl)-3,4-bis((*tert*-butyldimethylsilyl)oxy)tetrahydrofuran-2-yl)methyl)-*N*-(*tert*-butoxycarbonyl)-l-homocysteinate (6)**

Derivative **6** was synthesized from protected SAH analogue **1** (140 mg, 0.16 mmol) according to the same procedure as for compound **4**. FCC (3–35% of EtOAc in DCM) gave intermediate **6** (130 mg, 88 %)
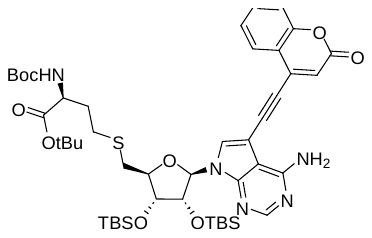
 as a yellow foam. **^1^H NMR** (400 MHz, DMSO-*d*_6_) δ 8.28 (s, 1H), 8.21 (s, 1H), 7.99 (dd, *J* = 8.1, 1.6 Hz, 1H), 7.70 (td, *J* = 7.5, 1.6 Hz, 1H), 7.49–7.43 (m, 2H), 7.18 (d, *J* = 7.8 Hz, 1H), 7.00 (s, 1H), 6.90 (bs, 2H), 6.15 (d, *J* = 7.2 Hz, 1H), 4.88 (dd, *J* = 7.3, 4.3 Hz, 1H), 4.23 (d, *J* = 4.6 Hz, 1H), 4.02 (t, *J* = 7.3 Hz, 1H), 3.98–3.89 (m, 1H), 3.07 (dd, *J* = 13.9, 8.4 Hz, 1H), 2.91 (dd, *J* = 13.9, 6.0 Hz, 1H), 2.70–2.43* (m, 2H), 1.95–1.76 (m, 2H), 1.37 (s, 9H), 1.36 (s, 9H), 0.94 (s, 9H), 0.68 (s, 9H), 0.17 (s, 3H), 0.13 (s, 3H), -0.09 (s, 3H), -0.38 (s, 3H). **^13^C NMR** (101 MHz, DMSO-*d*_6_) δ 171.6, 159.6, 157.7, 155.8, 153.7, 153.2, 151.0, 136.5, 132.9, 130.8, 126.7, 124.9, 118.0, 117.9, 117.5, 117.1, 102.0, 96.6, 93.9, 87.0, 85.6, 85.4, 80.6, 78.3, 74.4, 74.0, 53.7, 33.6, 31.2, 28.6, 28.4, 27.9, 25.8, 25.6, 17.9, 17.6, -4.3, -4.6, -4.7, -5.5. **LRMS** (ESI) [M + H]^+^ m/z: 935.5.

***tert*-Butyl *S*-(((2*S*,3*R*,4*R*,5*R*)-5-(4-amino-5-(2-(2-oxo-2*H*-chromen-4-yl)ethyl)-7*H*-pyrrolo[2,3-d]pyrimidin-7-yl)-3,4-bis((*tert*-butyldimejthylsilyl)oxy)tetrahydrofuran-2-yl)methyl)-*N*-(*tert*-butoxycarbonyl)-l-homocysteinate (7)**

Protected SAH analogue **7** was prepared from compound **6** (115 mg, 0.12 mmol) following the same methodology as for **5**. FCC (3–40% of EtOAc in DCM) furnished intermediate **7** (100 mg, 87%) as a yellow foam.
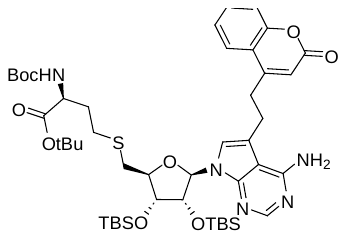
 **^1^H NMR** (400 MHz, DMSO-*d*_6_) δ 8.05 (s, 1H), 7.86 (dd, *J* = 8.0, 1.5 Hz, 1H), 7.63 (ddd, *J* = 8.5, 7.2, 1.5 Hz, 1H), 7.42 (dd, *J* = 8.4, 1.2 Hz, 1H), 7.38 (td, *J* = 7.0, 1.1 Hz, 1H), 7.29 (s, 1H), 7.15 (d, *J* = 7.9 Hz, 1H), 6.71 (bs, 2H), 6.56 (s, 1H), 6.09 (d, *J* = 7.4 Hz, 1H), 4.78 (dd, *J* = 7.2, 4.5 Hz, 1H), 4.19 (d, *J* = 4.5 Hz, 1H), 4.00–3.88 (m, 2H), 3.24–3.12 (m, 4H), 2.98 (dd, *J* = 13.7, 8.0 Hz, 1H), 2.80 (dd, *J* = 13.9, 5.9 Hz, 1H), 2.65–2.45* (m, 2H), 1.93–1.73 (m, 2H), 1.36 (s, 9H), 1.34 (s, 9iH), 0.93 (s, 9H), 0.67 (s, 8H), 0.15 (s, 3H), 0.12 (s, 3H), -0.13 (s, 3H), -0.39 (s, 3H). **^13^C NMR** (101 MHz, DMSO-*d*_6_) 171.60, 160.12, 157.83, 155.51, 153.24, 151.78, 151.63, 132.07, 124.94, 124.57, 119.40, 119.28, 116.90, 114.38, 113.72, 102.41, 86.32, 84.25, 80.52, 78.25, 74.68, 73.83, 53.46, 33.77, 31.20, 30.98, 28.44,28.32,27.74, 25.94, 5.66, 23.91, 17.93, 17.67, -4.44, -4.46, -4.58, -5.32. **LRMS** (ESI) [M + H]^+^ m/z: 940.4.

***S*-(((2*S*,3*S*,4*R*,5*R*)-5-(4-Amino-5-(2-(2-oxo-2*H*-chromen-4-yl)ethyl)-7*H*-pyrrolo[2,3-d]pyrimidin-7-yl)-3,4-dihydroxytetrahydrofuran-2-yl)methyl)-l-homocysteine (TO739)**

Final compound **TO739** was prepared from protected analogue **7** (85 mg, 0.09 mmol) using the same methodology as for **TO501**. RP-FCC (3–40% of MeOH in water) provided derivative **TO739** (44 mg, 87%) as a yellow solid.
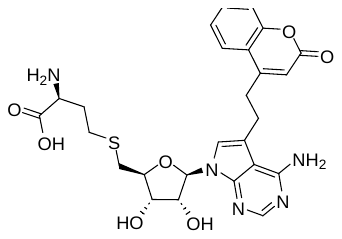
 **^1^H NMR** (400 MHz, DMSO-*d*_6_) δ 8.05 (s, 1H), 7.94 (dd, *J* = 8.0, 1.5 Hz, 1H), 7.62 (ddd, *J* = 8.5, 7.2, 1.5 Hz, 1H), 7.43–7.36 (m, 2H), 7.31 (s, 1H), 6.72 (bs, 2H), 6.63 (s, 1H), 6.07 (d, *J* = 5.8 Hz, 1H), 5.57 (bs, 2H), 4.44 (t, *J* = 5.6 Hz, 1H), 4.06 (t, *J* = 4.7, 1H), 3.95 (td, *J* = 6.3, 4.0 Hz, 1H), 3.34–3.29 (m, 1H), 3.18 (s, 4H), 2.88 (dd, *J* = 13.7, 6.1 Hz, 1H), 2.72 (dd, *J* = 13.6, 6.7 Hz, 1H), 2.64 (t, *J* = 7.8 Hz, 2H), 2.07–1.94 (m, 1H), 1.90–1.77 (m, 1H). **^13^C NMR** (101 MHz, DMSO-*d*_6_) δ 170.1, 160.3, 157.9, 155.9, 153.2, 151.9, 151.4, 132.1, 125.4, 124.7, 119.4, 118.9, 116.8, 114.5, 113.6, 102.2, 86.7, 82.7, 73.3, 72.8, 53.2, 34.3, 31.6, 30.7, 28.4, 24.1. **HRMS** (ESI) m/z: [M + H]^+^ (C_26_H_29_N_5_O_7_S) calculated 556.186, found 556.186.

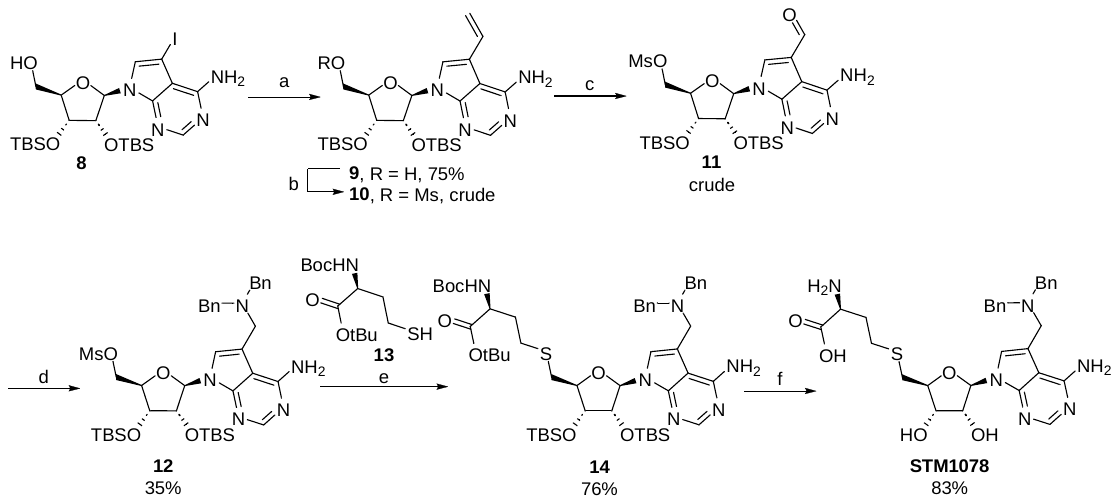

**SI Scheme 1** – Reagents and conditions: (a) tributyl(vinyl)stannane, Pd(PPh_3_)_4_, DMF, 110 °C, 4 h; (b) MsCl, TEA, DCM, RT, 30 min; (c) (i) OsO_4_, *N*-Methylmorpholine *N*-oxide, THF/water (2:1), 0 °C to RT, 3 h; (ii) NaIO_4_, THF/water (2:1), RT, overnight; (d) dibenzylamine, NaBH(OAc)_3_, AcOH, DCM, RT, 3 d (e) **13**, *t-*BuOK, NMP, 0 °C to RT, 1 d; (f) 90% TFA (aq.), RT, overnight.

**((2*R*,3*R*,4*S*,5*R*)-5-(4-Amino-5-vinyl-7*H*-pyrrolo[2,3-*d*]pyrimidin-7-yl)-3,4-bis((*tert*-butyldimethylsilyl)oxy)tetrahydrofuran-2-yl)methanol (9)**

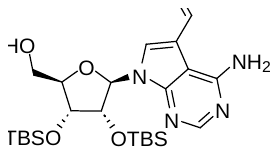
Starting material **8** (621 mg, 1.00 mmol), prepared according to a reported procedure,[2] and Pd(PPh_3_)_4_ (116 mg, 0.10 mmol, 0.10 eq) were dissolved in anhydrous DMF (10 mL) and the mixture was degassed and filled with argon. Vinyltributylstannane (1.2 mL, 4.00 mmol, 4.0 eq) was added and the reaction mixture was stirred at 110 °C for 4 h. The solvent was removed under reduced pressure, and the residue was purified by FCC (10–30% of EtOAc/EtOH (4:1) in DCM). Impure fractions were collected separately and subjected to repeated chromatography under the same conditions, providing compound **9** (390 mg, 75%) as a yellowish solid. **^1^H NMR** (401 MHz, DMSO*-d*_6_) δ 8.04 (s, 1H), 7.69 (s, 1H), 7.11 (dd, *J* = 17.2, 10.9 Hz, 1H), 6.74 (s, 2H), 6.04 (d, *J* = 7.3 Hz, 1H), 5.58–5.49 (m, 2H), 5.12 (dd, *J* = 10.9, 1.8 Hz, 1H), 4.70 (dd, *J* = 7.3, 4.6 Hz, 1H), 4.28–4.22 (m, 1H), 3.97–3.90 (m,1H), 3.74–3.64 (m, 1H), 3.57 (ddd, *J* = 12.0, 6.8, 3.2 Hz, 1H), 0.92 (s, 9H), 0.68 (s, 9H), 0.12 (s, 3H), 0.11 (s, 3H), -0.16 (s, 3H), -0.43 (s, 3H). **^13^C NMR** (101 MHz, DMSO*-d*_6_) δ 157.8, 151.6, 150.9, 129.2, 119.6, 114.4, 113.3, 101.2, 87.0, 86.6, 74.8, 73.3, 61.7, 26.0, 25.7, 18.0, 17.7, -4.5, -4.5, -4.6, -5.4. **LRMS** (ESI) [M + H]^+^ m/z: 521.1.

**((2*R*,3*R*,4*S*,5*R*)-5-(4-Amino-5-formyl-7*H*-pyrrolo[2,3-*d*]pyrimidin-7-yl)-3,4-bis((*tert*-butyldimethylsilyl)oxy)tetrahydrofuran-2-yl)methyl methanesulfonate (10)**

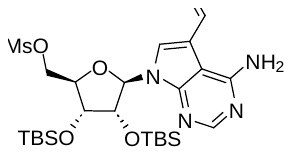
Starting material **9** (260 mg, 0.50 mmol) was dissolved in DCM (5.0 mL, dried over Na_2_SO_4_). Triethylamine (97 µL, 0.70 mmol, 1.4 eq) was added, followed by methanesulfonyl chloride (46 µL, 0.60 mmol, 1.2 eq). The reaction mixture was stirred at RT for 30 min. The mixture was then diluted with DCM, washed with water and brine, dried over Na_2_SO_4_, and concentrated under reduced pressure, providing the crude mesylate **10** (316 mg), which was used in the next step without further purification. **^1^H NMR** (401 MHz, DMSO*-d*_6_) δ 8.06 (s, 1H), 7.66 (s, 1H), 7.12 (dd, *J* = 17.2, 10.9 Hz, 1H), 6.73 (s, 2H), 6.17 (d, *J* = 7.2 Hz, 1H), 5.58 (dd, *J* = 17.2, 1.7 Hz, 1H), 5.13 (dd, *J* = 10.9, 1.8 Hz, 1H), 4.71 (dd, *J* = 7.2, 4.5 Hz, 1H), 4.55–4.47 (m, 2H), 4.29 (dd, *J* = 4.5, 1.4 Hz, 1H), 4.13 (td, *J* = 5.1, 1.4 Hz, 1H), 3.21 (s, 3H), 0.93 (s, 9H), 0.67 (s, 9H), 0.15 (s, 3H), 0.12 (s, 3H), -0.11 (s, 3H), -0.36 (s, 3H). **^13^C NMR** (101 MHz, DMSO*-d*_6_) δ 157.8, 152.0, 151.6, 129.1, 118.6, 115.0, 113.4, 100.9, 86.0, 82.5, 74.4, 72.6, 69.6, 37.0, 25.9, 25.6, 17.9, 17.7, -4.5, -4.6, -4.7, -5.4. **LRMS** (ESI) [M + H]^+^ m/z: 599.2.

**((2*R*,3*R*,4*S*,5*R*)-5-(4-Amino-5-formyl-7*H*-pyrrolo[2,3-*d*]pyrimidin-7-yl)-3,4-bis((*tert*-butyldimethylsilyl)oxy)tetrahydrofuran-2-yl)methyl methanesulfonate (11)**

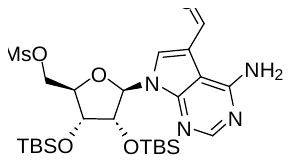
Mesylate **10** (299 mg, 0.50 mmol) was dissolved in a 2:1 mixture of THF (4.0 mL) and water (2.0 mL), and the solution was cooled to 0 °C. A 4% aqueous solution of OsO_4_ (159 µL, 0.025 mmol, 0.05 eq) and a 50 wt% aqueous *N*-methylmorpholine *N*-oxide solution (114 µL, 0.549 mmol, 1.1 eq) were added and the reaction mixture was stirred for 3 h, while gradually warming to RT. After complete dihydroxylation of the vinyl group, sodium periodate (160 mg, 0.749 mmol, 1.5 eq) was added and stirring was continued overnight. The reaction mixture was diluted with EtOAc and washed successively with aqueous Na_2_S_2_O_3_ solution, water, and brine, then dried over Na_2_SO_4_. Concentration of the organic phase under reduced pressure provided crude aldehyde **11** (332 mg), which was used directly in the next step. ^1^H NMR (401 MHz, DMSO*-d*_6_) δ 9.72 (s, 1H), 8.53 (s, 1H), 8.21 (s, 1H), 7.63 (s, 2H), 6.15 (d, *J* = 6.5 Hz, 1H), 4.81 (dd, *J* = 6.5, 4.5 Hz, 1H), 4.58 (dd, *J* = 11.2, 4.8 Hz, 1H), 4.53 (dd, *J* = 11.2, 6.0 Hz, 1H), 4.34 (dd, *J* = 4.4, 2.3 Hz, 1H), 4.22–4.17 (m, 1H), 3.23 (s, 3H), 0.92 (s, 9H), 0.71 (s, 9H), 0.15 (s, 3H), 0.12 (s, 3H), -0.06 (s, 3H), -0.33 (s, 3H). The ^13^C NMR spectrum of compound **114** was not recorded. **LRMS** (ESI) [M + H]^+^ m/z: 601.2.

**((2*R*,3*R*,4*S*,5*R*)-5-(4-Amino-5-((dibenzylamino)methyl)-7*H*-pyrrolo[2,3-*d*]pyrimidin-7-yl)-3,4-bis((*tert*-butyldimethylsilyl)oxy)tetrahydrofuran-2-yl)methyl methanesulfonate (12)**

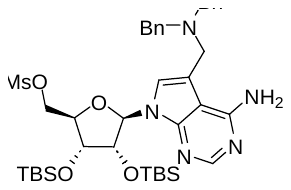
Aldehyde **11** (100 mg, 0.166 mmol) and dibenzylamine (35 µL, 0.183 mmol, 1.1 eq) were dissolved in DCM (1.7 mL), followed by the addition of NaBH(OAc)_3_ (53 mg, 0.25 mmol, 1.5 eq) and AcOH (10 µL, 0.166 mmol, 1.0 eq). The reaction mixture was stirred at RT for 2 days, then an additional portion of dibenzylamine (95 µL, 0.50 mmol, 3.0 eq) was added, and stirring was continued for 1 day at the same temperature. Volatiles were removed under reduced pressure, and the residue was purified by RP-FCC (40–100% of ACN in water), furnishing the amine **12** (45 mg, 0.058 mmol, 35%). ^1^H NMR (401 MHz, DMSO*-d*_6_) δ 8.05 (s, 1H), 7.40 (s, 1H), 7.38–7.23 (m, 12zH), 6.10 (d, *J* = 7.3 Hz, 1H), 4.70 (dd, *J* = 7.3, 4.6 Hz, 1H), 4.55–4.44 (m, 2H), 4.31–4.25 (m, 1H), 4.16–4.07 (m, 1H), 3.68–3.54 (m, 4H), 3.53–3.40 (m, 2H), 3.21 (s, 3H), 0.92 (s, 9H), 0.60 (s, 9H), 0.13 (s, 3H), 0.11 (s, 3H), -0.15 (s, 3H), -0.43 (s, 3H). The ^13^C NMR spectrum of the compound was not included due to poor data quality caused by its instability. **LRMS** (ESI) [M + H]^+^ m/z: 782.3.

***tert*-Butyl *S*-(((2*R*,3*R*,4*S*,5*R*)-5-(4-amino-5-((dibenzylamino)methyl)-7*H*-pyrrolo[2,3-*d*]pyrimidin-7-yl)-3,4-bis((*tert*-butyldimethylsilyl)oxy)tetrahydrofuran-2-yl)methyl)-*N*-(*tert*-butoxycarbonyl)-l-homocysteinate (14)**

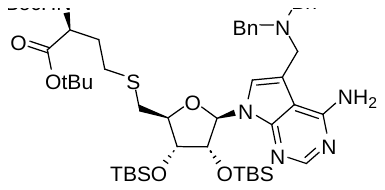
To an ice-cold solution of compound **12** (38 mg, 0.049 mmol) in NMP (1.0 mL) was added a solution of protected homocysteine **13** (32 mg, 0.097 mmol, 2.0 eq), prepared according to a published procedure,[3] and *t*-BuOK (11 mg, 0.097 mmol, 2.0 eq) in NMP (0.5 mL), pre-mixed for 5 min at 0 °C. The reaction mixture was gradually warmed to RT and stirred for 24 h. The mixture was diluted with EtOAc, washed three times with water and once with brine, and dried over Na_2_SO_4_. After filtration and concentration under reduced pressure, the residue was purified by RP-FCC (30–100% of ACN in water), providing protected SAH analog **14** (36 mg, 76%). **^1^H NMR** (401 MHz, DMSO*-d*_6_) δ 8.03 (s, 1H), 7.39 (s, 1H), 7.38–7.25 (m, 12H), 7.16 (d, *J* = 7.9 Hz, 1H), 6.06 (d, *J* = 7.4 Hz, 1H), 4.76 (dd, *J* = 7.3, 4.4 Hz, 1H), 4.17 (d, *J* = 4.5 Hz, 1H), 3.93 (dt, *J* = 14.1, 7.4 Hz, 2H), 3.69–3.51 (m, 4H), 3.50–3.42 (m, 2H), 2.99 (dd, *J* = 13.8, 8.3 Hz, 1H), 2.82 (dd, *J* = 13.8, 5.8 Hz, 1H), 2.64–2.49 (m, 2H), 1.87–1.78 (m, 2H), 0.92 (s, 9H), 0.59 (s, 9H), 0.14 (s, 3H), 0.11 (s, 3H), -0.16 (s, 3H), -0.46 (s, 3H). **^13^C NMR** (101 MHz, DMSO*-d*_6_) δ 158.1, 155.8, 152.3*, 151.8, 137.5, 129.6, 128.6, 127.5, 121.7*, 112.7, 102.5, 86.2, 84.5, 80.5, 78.3, 74.7, 74.0, 57.3, 53.5, 50.3, 33.7, 31.2, 28.42*, 28.36, 27.8, 26.0, 25.6, 17.9, 17.6, -4.41, -4.44, -4.5, -5.4. **LRMS** (ESI) [M + H]^+^ m/z: 977.4.

***S*-(((2*S*,3*S*,4*R*,5*R*)-5-(4-Amino-5-((dibenzylamino)methyl)-7*H*-pyrrolo[2,3-*d*]pyrimidin-7-yl)-3,4-dihydroxytetrahydrofuran-2-yl)methyl)-l-homocysteine (STM1078)**

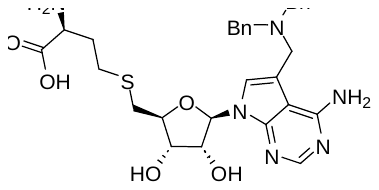
Compound **14** (36 mg, 0.037 mmol) was dissolved in 90% aqueous TFA (3.0 mL), and the reaction mixture was stirred at RT overnight. Volatiles were removed under reduced pressure, the residue was adsorbed onto silica and purified by RP-FCC (25–100% of ACN in water, 0.1% FA). The compound was freeze-dried to remove residual formic acid, providing SAH analog **STM1078** (18 mg, 83%) as a white solid. **^1^H NMR** (401 MHz, DMSO*-d*_6_) δ 8.04 (s, 1H), 7.38–7.24 (m, 13H), 6.02 (d, *J* = 5.8 Hz, 1H), 4.41 (t, *J* = 5.5 Hz, 1H), 4.07–4.01 (m, 1H), 3.99–3.90 (m, 1H), 3.66–3.47 (m, 6H), 3.40 (t, *J* = 6.4 Hz, 1H), 2.88 (dd, *J* = 13.6, 6.1 Hz, 1H), 2.74 (dd, *J* = 13.6, 6.5 Hz, 1H), 2.63 (t, *J* = 7.5 Hz, 2H), 2.07–1.92 (m, 1H), 1.90–1.75 (m, 1H). **^13^C NMR** (101 MHz, DMSO*-d*_6_) δ 170.0, 158.0, 152.3, 151.5, 137.5, 129.6, 128.6, 127.5, 121.5, 112.5, 102.3, 86.7, 82.8, 73.3, 72.7, 57.4, 52.8, 50.5, 34.2, 31.3, 28.2. **HRMS** (ESI) m/z: [M + H]^+^ (C_30_H_37_N_6_O_5_S) calculated 593.2541, found 593.2544.

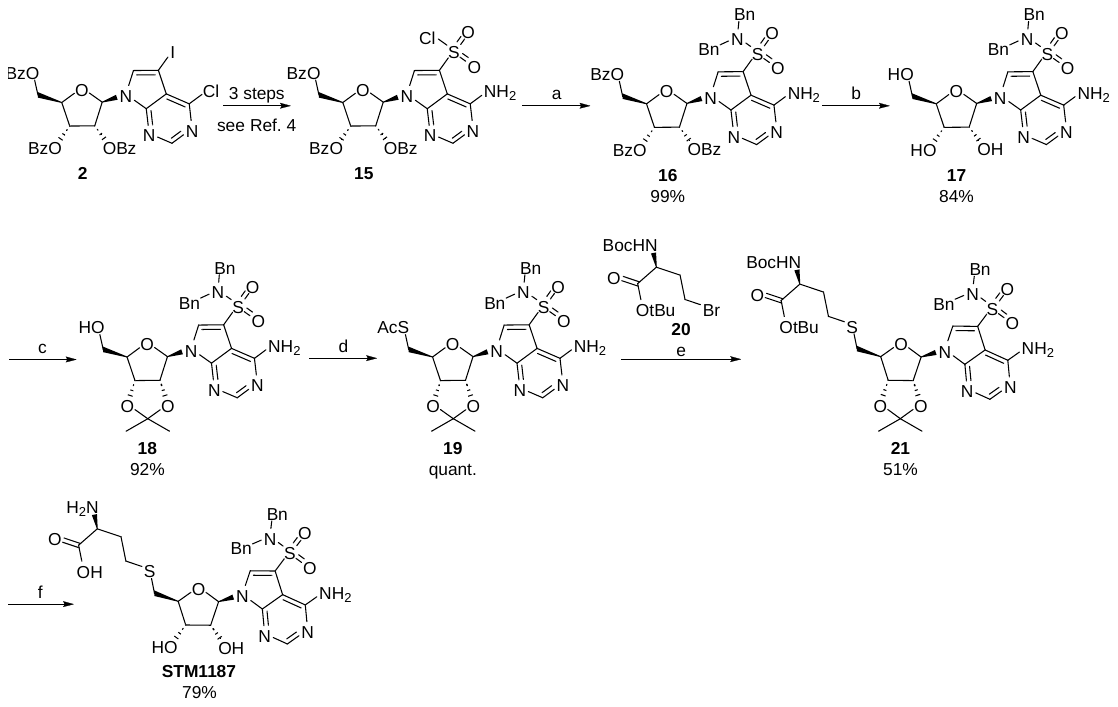

**SI Scheme 2** – Reagents and conditions: (a) dibenzylamine, DCM, RT, overnight; (b) 7 M NH_3_ in MeOH, RT, 20 h; (c) 2,2-dimethoxypropane, acetone, RT, 1 h; (d) (i) AcSH, PPh_3_, DIAD, THF, 0 °C to RT, 90 min, (ii) AcOH, H_2_O, THF, 65 °C, 5 h; (e) **20**, K_2_CO_3_, PPh_3_, MeOH, 0 °C to RT, 1 d; (f) 90% TFA (aq.), RT, 3 h.

**(2*R*,3*R*,4*R*,5*R*)-2-(4-Amino-5-(chlorosulfonyl)-7*H*-pyrrolo[2,3-*d*]pyrimidin-7-yl)-5-((benzoyloxy)methyl)tetrahydrofuran-3,4-diyl dibenzoate (15)**

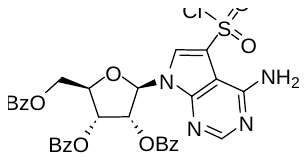
Compound **15** was prepared from intermediate **2** in three steps following our previously reported procedure.[4] Intermediate **2** was synthesized as described previously.[5]

**(2*R*,3*R*,4*R*,5*R*)-2-(4-Amino-5-(*N*,*N*-dibenzylsulfamoyl)-7*H*-pyrrolo[2,3-*d*]pyrimidin-7-yl)-5-((benzoyloxy)methyl)tetrahydrofuran-3,4-diyl dibenzoate (16)**

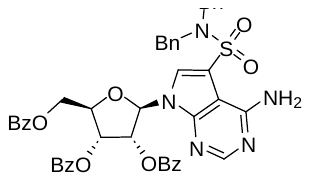
Sulfonyl chloride **15** (241 mg, 0.36 mmol) was dissolved in anhydrous DCM (5.1 mL). Dibenzylamine (151 µL, 0.78 mmol, 2.2 eq) was added at 0 °C, and the reaction was stirred for 30 minutes before the bath was removed, and the stirring was continued overnight. The resulting suspension was diluted with MeOH, affording a clear solution, which was adsorbed onto silica. FCC (10–40% of EtOAc/EtOH (4:1) in cyclohexane) yielded **16** (296 mg, 99%) as a colorless solid. **^1^H NMR** (401 MHz, CDCl_3_) δ 8.28 (s, 1H), 8.12*–*8.05 (m, 2H), 8.04*–*7.98 (m, 2H), 7.94*–*7.87 (m, 2H), 7.62 (s, 1H), 7.61*–*7.47 (m, 3H), 7.44*–*7.33 (m, 6H), 7.23*–*7.12 (m, 6H), 7.09*–*7.00 (m, 4H), 6.58 (d, *J* = 5.3 Hz, 1H), 6.17*–*6.07 (m, 2H), 4.83 (dd, *J* = 11.8. 3.1 Hz, 1H), 4.81*–*4.77 (m, 1H), 4.71 (dd, *J* = 11.8, 3.9 Hz, 1H), 4.26 (s, 4H). **^13^C NMR** (101 MHz, CDCl_3_) δ 166.3, 165.5, 165.2, 156.8, 153.2*, 151.3, 135.4, 133.9, 133.9, 133.7, 130.0, 130.0, 129.8, 129.3, 128.9, 128.7, 128.7, 128.6, 128.6, 127.9, 127.1, 116.4, 99.8, 87.1, 81.0, 74.2, 71.7, 63.9, 51.1. **HRMS** (ESI) m/z: [M+H]^+^ (C_46_H_40_N_5_O_9_S) calculated 838.2541, found 838.2538.

**4-Amino-*N*,*N*-dibenzyl-7-((2*R*,3*R*,4*S*,5*R*)-3,4-dihydroxy-5-(hydroxymethyl)tetrahydrofuran-2-yl)-7*H*-pyrrolo[2,3*-d*]pyrimidine-5-sulfonamide (17)**

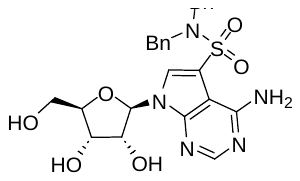
Starting protected nucleoside analog **16** (160 mg, 0.19 mmol) was dissolved in 7 M methanolic ammonia (5.3 mL) and stirred at RT for 20 h. After this time, the volatiles were removed under reduced pressure, and the residue was adsorbed onto silica. Purification by FCC (2–20% of MeOH in DCM) afforded **17** (84 mg, 84%). **^1^H NMR** (401 MHz, DMSO-*d*_6_) δ 8.42 (s, 1H), 8.24 (s, 1H), 7.25*–*7.15 (m, 6H), 7.13*–*7.05 (m, 4H), 6.12 (d, *J* = 5.0 Hz, 1H), 5.49 (d, *J* = 5.9 Hz, 1H), 5.28 (t, *J* = 5.3 Hz, 1H), 5.17 (d, *J* = 5.1 Hz, 1H), 4.43*–*4.35 (m, 1H), 4.34*–*4.22 (m, 4H), 4.18*–*4.08 (m, 1H), 3.99*–*3.92 (m, 1H), 3.76*–*3.67 (m, 1H), 3.62*–*3.53 (m, 1H). **^13^C NMR** (101 MHz, DMSO-*d*_6_) δ 157.1, 153.4, 150.8, 136.4, 128.7, 128.4, 128.3, 127.6, 112.5, 98.7, 88.2, 85.3, 74.5, 70.0, 61.0, 51.6. **HRMS** (ESI) m/z: [M+H]^+^ (C_25_H_28_N_5_O_6_S) calculated 526.1755, found 526.1751.

**4-Amino-*N*,*N*-dibenzyl-7-((3a*R*,4*R*,6*R*,6a*R*)-6-(hydroxymethyl)-2,2-dimethyltetrahydrofuro[3,4-*d*][1,3]dioxol-4-yl)-7*H*-pyrrolo[2,3-*d*]pyrimidine-5-sulfonamide (18)**

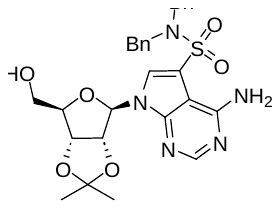
Nucleoside analog **17** (70 mg, 0.133 mmol) and TsOH·H_2_O (25 mg, 0.133 mmol, 1.0 eq) were dissolved in a mixture of acetone (1.6 mL) and 2,2-dimethoxypropane (0.40 mL), and the reaction mixture was stirred at RT for 1 h. After this time, MeOH (0.20 mL) was added, and the mixture was stirred overnight. Volatiles were removed under reduced pressure, and the residue was purified by RP-FCC (30–100% of ACN in water), furnishing compound **18** (69 mg, 0.12 mmol, 92%) as an off-white solid. **^1^H NMR** (400 MHz, CDCl_3_) δ 8.30 (s, 1H), 7.33 (s, 1H), 7.25–7.21 (m, 6H), 7.15–7.10 (m, 4H), 6.09 (d, *J* = 10.3 Hz, 1H), 5.63 (d, *J* = 5.0 Hz, 1H), 5.26–5.19 (m, 1H), 5.10 (dd, *J* = 6.0, 1.3 Hz, 1H), 4.54–4.48 (m, 1H), 4.42–4.29 (m, 4H), 4.01–3.92 (m, 1H), 3.85–3.76 (m, 1H), 1.64 (s, 3H), 1.38 (s, 3H). **^13^C NMR** (101 MHz, CDCl_3_) δ 157.6, 153.3, 149.5, 135.4, 130.0, 128.7, 128.6, 128.1, 114.4, 114.2, 101.0, 96.8, 86.1, 82.8, 81.6, 63.5, 51.1, 27.8, 25.4. **LRMS** (ESI) [M + H]^+^ m/z: 566.1.

***S*-(((3a*S*,4*S*,6*R*,6a*R*)-6-(4-Amino-5-(*N*,*N*-dibenzylsulfamoyl)-7*H*-pyrrolo[2,3-*d*]pyrimidin-7-yl)-2,2-dimethyltetrahydrofuro[3,4-*d*][1,3]dioxol-4-yl)methyl) ethanethioate (19)**

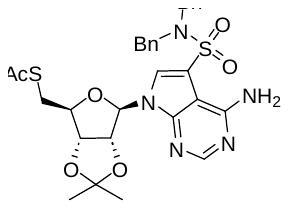
Starting material **18** (68 mg, 0.12 mmol) and PPh_3_ (79 mg, 0.30 mmol, 2.5 eq) were dissolved in anhydrous THF (1.2 mL) and the solution was cooled to 0 °C in an ice bath. DIAD (59 µL, 0.30 mmol, 2.5 eq) was added and the reaction mixture was stirred at 0 °C for 30 min. Thioacetic acid (19 µL, 0.27 mmol, 2.2 eq) was then added and stirring was continued for 1 h while allowing the temperature to rise to RT. Water (200 µL) and acetic acid (41 µL) were added and the mixture was stirred at 65 °C for 5 h to hydrolyze the formed N6-iminophosphorane intermediate. Volatiles were removed under reduced pressure, and the residue was purified by RP-FCC (30–100% of ACN in water), furnishing compound **19** (75 mg, quantitative). **^1^H NMR** (400 MHz, CDCl_3_) δ 8.37 (s, 1H), 7.47 (s, 1H), 7.25–7.18 (m, 6H), 7.11 (s, 4H), 6.10–6.04 (m, 1H), 5.20–5.14 (m, 1H), 4.82 (dd, *J* = 6.2, 3.5 Hz, 1H), 4.37 (s, 4H), 4.34–4.26 (m, 1H), 3.28 (dd, *J* = 13.8, 6.9 Hz, 1H), 3.18 (dd, *J* = 13.8, 6.1 Hz, 1H), 2.34 (s, 3H), 1.60 (s, 3H), 1.38 (s, 3H). **^13^C NMR** (101 MHz, CDCl_3_) δ 194.6, 157.2, 154.0, 150.8, 135.4, 128.7, 128.6, 128.2, 128.0, 114.96, 114.95, 99.9, 91.5, 85.7, 84.9, 83.3, 51.0, 31.3, 30.7, 27.3, 25.5. **LRMS** (ESI) [M + H]^+^ m/z: 624.2.

***tert*-Butyl *S*-(((3a*S*,4*S*,6*R*,6a*R*)-6-(4-amino-5-(*N*,*N*-dibenzylsulfamoyl)-7*H*-pyrrolo[2,3-*d*]pyrimidin-7-yl)-2,2-dimethyltetrahydrofuro[3,4-*d*][1,3]dioxol-4-yl)methyl)-*N*-(*tert*-butoxycarbonyl)-l-homocysteinate (21)**

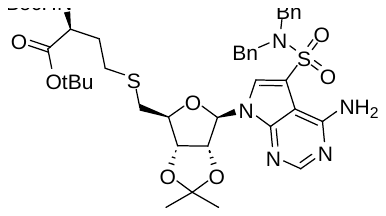
A solution of thioester **19** (74 mg, 0.12 mmol) in anhydrous MeOH (2.1 mL) was deoxygenated by repeated evacuation–refilling with argon and cooled in an ice bath. PPh_3_ (4 mg, 0.015 mmol, 0.13 eq), bromide **20** (60 mg, 0.18 mmol, 1.5 eq), prepared according to a literature procedure,[6] and finely ground K_2_CO_3_ (34 mg, 0.25 mmol, 2.1 eq) were added, and the flask was again evacuated and filled with argon. After stirring at 0 °C for 1 h the reaction mixture was allowed to warm to RT and stirred for 10 h. An additional portion of **20** (60 mg, 0.18 mmol, 1.5 eq) was added and stirring was continued overnight. The reaction mixture was adsorbed onto silica and purified by RP-FCC (20–100% of ACN in water), providing protected SAH analog **21** (51 mg, 51%). **^1^H NMR** (401 MHz, CDCl_3_) δ 8.35 (s, 1H), 7.50 (s, 1H), 7.25–7.19 (m, 6H), 7.16–7.06 (m, 4H), 6.07 (d, *J* = 2.3 Hz, 1H), 5.18–5.12 (m, 1H), 5.09 (d, *J* = 7.7 Hz, 1H), 4.95–4.90 (m, 1H), 4.34–4.30 (m, 1H), 4.28–4.20 (m, 1H), 2.84 (dd, *J* = 13.6, 7.1 Hz, 1H), 2.78 (dd, *J* = 13.6, 5.7 Hz, 1H), 2.62–2.54 (m, 2H), 2.11–1.98 (m, 1H), 1.90–1.79 (m, 1H), 1.62 (s, 3H), 1.44 (s, 9H), 1.42 (s, 9H), 1.39 (s, 3H). **^13^C NMR** (101 MHz, CDCl_3_) δ 171.3, 157.2, 154.0, 150.8, 135.5, 128.7, 128.6, 128.3, 128.0, 114.94, 114.91*, 99.9, 91.5, 86.0, 84.8, 83.5, 82.4, 79.9*, 53.4, 51.1, 34.6, 33.2, 28.8, 28.5, 28.1, 27.3, 25.5. **LRMS** (ESI) [M + H]^+^ m/z: 839.2.

***S*-(((2*S*,3*S*,4*R*,5*R*)-5-(4-Amino-5-(*N*,*N*-dibenzylsulfamoyl)-7*H*-pyrrolo[2,3-*d*]pyrimidin-7-yl)-3,4-dihydroxytetrahydrofuran-2-yl)methyl)-l-homocysteine (STM1187)**

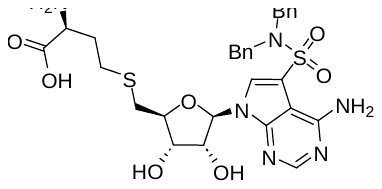
Compound **21** (48 mg, 0.057 mmol) was dissolved in 90% aqueous TFA (1.5 mL) and stirred at RT for 6 h. After complete deprotection, volatiles were removed under reduced pressure and the residue was purified by RP-FCC (10–70% of ACN in water, 0.1% FA), furnishing compound **STM1187** (29 mg, 79%) as a clear solid. **^1^H NMR** (401 MHz, DMSO*-d*_6_) δ 8.31 (s, 1H), 8.25 (s, 1H), 7.62 (bs, 3H), 7.27–7.15 (m, 6H), 7.14–7.07 (m, 4H), 6.14 (d, *J* = 5.9 Hz, 1H), 5.59 (bs, 1H), 4.61 (t, *J* = 5.5 Hz, 1H), 4.37–4.24 (m, 4H), 4.12–4.06 (m, 1H), 4.06–3.99 (m, 1H), 3.31–3.23 (m, 1H), 2.95 (dd, *J* = 13.8, 6.4 Hz, 1H), 2.80 (dd, *J* = 13.7, 6.5 Hz, 1H), 2.63 (t, *J* = 7.7 Hz, 2H), 2.06–1.93 (m, 1H), 1.88–1.75 (m, 1H). **^13^C NMR** (101 MHz, DMSO*-d*_6_) δ 169.7, 157.1, 153.5, 151.3, 136.4, 128.8, 128.4, 128.3, 127.6, 113.1, 98.6, 87.7, 83.5, 73.2, 72.5, 53.2, 51.6, 33.8, 31.6, 28.3. **HRMS** (ESI) m/z: [M + H]^+^ (C_29_H_35_N_6_O_7_S_2_) calculated 643.2003, found 643.2005.

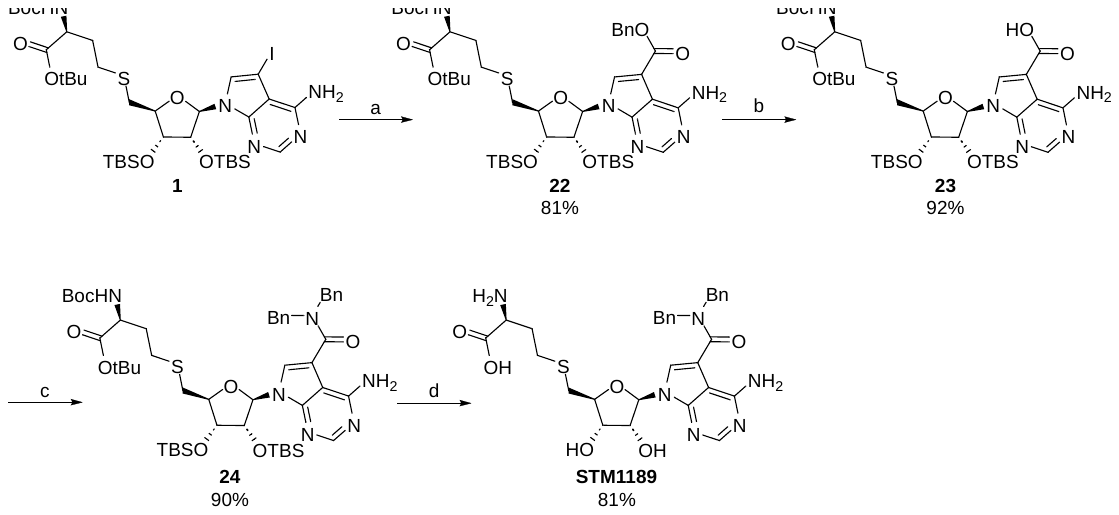

**SI Scheme 3** – Reagents and conditions: (a) Mo(CO)_6_, Pd(dppf)Cl_2_·DCM, DBU, BnOH, 80 °C, 30 min; (b) 10% Pd/C, H_2_, EtOAc, RT, overnight; (c) dibenzylamine, HATU, DIPEA, DMF, RT, overnight; (d) 90% TFA (aq.), RT, 3 h.

**Benzyl 4-amino-7-((2*R*,3*R*,4*R*,5*S*)-5-((((*S*)-4-(*tert*-butoxy)-3-((*tert*-butoxycarbonyl)amino)-4-oxobutyl)thio)methyl)-3,4-bis((*tert*-butyldimethylsilyl)oxy)tetrahydrofuran-2-yl)-7*H*-pyrrolo[2,3-*d*]pyrimidine-5-carboxylate (22)**

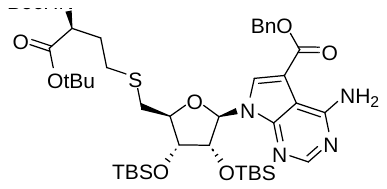
Starting material **1** (200 mg, 0.22 mmol) was suspended in anhydrous BnOH (2.2 mL) together with Mo(CO)_6_ (59 µL, 0.22 mmol, 1.0 eq) and Pd(dppf)·DCM (18.3 mg, 0.022 mmol, 0.10 eq). DBU (100 µL, 0.67 mmol, 3.0 eq) was added and the reaction mixture was stirred at 80 °C for 30 min. The reaction mixture was transferred into a larger flask and BnOH was removed by repeated azeotropic co-distillation with water. The residue was adsorbed onto silica and purified by RP-FCC (30–100% of ACN in water), providing compound **22** (163 mg, 81%) as a light brown oil. **^1^H NMR** (400 MHz, DMSO*-d*_6_) δ 8.33 (s, 1H), 8.16 (s, 1H), 7.79 (s, 1H), 7.50 (s, 1H), 7.48–7.43 (m, 2H), 7.43–7.31 (m, 3H), 7.16 (d, *J* = 7.9 Hz, 1H), 6.15 (d, *J* = 7.1 Hz, 1H), 5.41 (d, *J* = 12.9 Hz, 1H), 5.35 (d, *J* = 12.9 Hz, 1H), 4.99–4.93 (m, 1H), 4.20 (m, 1H), 4.03–3.99 (m, 1H), 3.93–3.86 (m, 1H), 3.09 (dd, *J* = 13.8, 8.1 Hz, 1H), 2.87 (dd, *J* = 14.0, 5.8 Hz, 1H), 2.62–2.44* (m, 2H), 1.88–1.77 (m, 2H), 1.37 (s, 9H), 1.35 (s, 9H), 0.92 (s, 9H), 0.67 (s, 9H), 0.15 (s, 3H), 0.11 (s, 3H), -0.10 (s, 3H), -0.39 (s, 3H). **^13^C NMR** (101 MHz, DMSO*-d*_6_) δ 171.6, 164.8, 157.7, 155.7, 153.6, 151.9, 136.5, 130.8, 128.7, 128.2, 127.7, 106.7, 100.8*, 87.2, 84.9, 80.5, 78.3, 74.4, 73.8, 65.9, 53.5, 33.3, 31.2, 28.5, 28.3, 27.8, 25.9, 25.6, 17.9, 17.6, -4.44, -4.46, -4.53, -5.4. **LRMS** (ESI) [M + H]^+^ m/z: 902.1.

**4-Amino-7-((2*R*,3*R*,4*R*,5*S*)-5-((((*S*)-4-(*tert*-butoxy)-3-((*tert*-butoxycarbonyl)amino)-4-oxobutyl)thio)methyl)-3,4-bis((*tert*-butyldimethylsilyl)oxy)tetrahydrofuran-2-yl)-7*H*-pyrrolo[2,3-*d*]pyrimidine-5-carboxylic acid (23)**

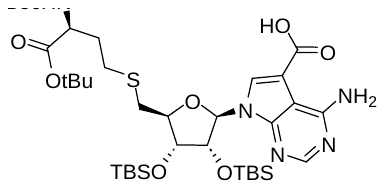
Starting material **22** (212 mg, 0.235 mmol) was dissolved in EtOAc (2.4 mL). The reaction flask was repeatedly evacuated and filled with argon before the addition of 10% Pd/C (21 mg, 10 w%). Hydrogen was then introduced by repeated evacuation–refilling cycles, and the reaction mixture was stirred overnight under a hydrogen atmosphere. The catalyst was removed by filtration through a pad of celite, which was washed thoroughly with EtOAc. Concentration under reduced pressure furnished compound **23** (175 mg, 92%). **^1^H NMR** (400 MHz, DMSO*-d*_6_) δ 12.98 (s, 1H), 8.16 (s, 1H), 8.13 (s, 1H), 7.41 (bs, 1H), 7.16 (d, *J* = 7.9 Hz, 1H), 6.15 (d, *J* = 7.3 Hz, 1H), 4.89 (dd, *J* = 7.3, 4.4 Hz, 1H), 4.19 (d, *J* = 4.5 Hz, 1H), 4.07–3.80 (m, 2H), 3.08 (dd, *J* = 13.9, 8.4 Hz, 1H), 2.86 (dd, *J* = 13.9, 5.9 Hz, 1H), 2.61–2.45* (m, 2H), 1.90–1.75 (m, 2H), 1.41–1.32 (m, 18H), 0.93 (s, 9H), 0.67 (s, 9H), 0.15 (s, 3H), 0.12 (s, 3H), -0.10 (s, 3H), -0.41 (s, 3H). **^13^C NMR** (101 MHz, DMSO*-d*_6_) δ 171.6, 166.7, 157.9, 155.7, 151.9, 130.1, 100.9, 86.7, 84.9, 80.5, 78.3, 74.6, 74.0, 53.5, 33.4, 31.2, 28.4, 28.3, 27.8, 26.0, 25.6, 17.9, 17.6, -4.4, -4.6, -5.4. **LRMS** (ESI) [M + H]^+^ m/z: 812.3.

***tert*-Butyl *S*-(((2*R*,3*R*,4*S*,5*R*)-5-(4-amino-5-(dibenzylcarbamoyl)-7*H*-pyrrolo[2,3-*d*]pyrimidin-7-yl)-3,4-bis((*tert*-butyldimethylsilyl)oxy)tetrahydrofuran-2-yl)methyl)-*N*-(*tert*-butoxycarbonyl)-l-homocysteinate (24)**

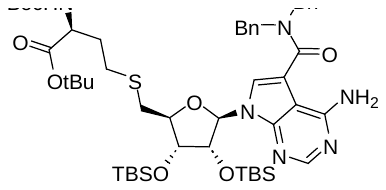
To a stirred solution of acid **23** (81 mg, 0.10 mmol) and HATU (42 mg, 0.11 mmol, 1.1 eq) in anhydrous DMF (1.0 mL) was added dibenzylamine (29 µL, 0.15 mmol, 1.5 eq), followed by DIPEA (35 µL, 0.20 mmol, 2.0 eq). The resulting mixture was stirred at RT for 3 h. The solvent was removed under reduced pressure, and the residue was purified by FCC (5–25% of EtOAc/EtOH (4:1) in cyclohexane), giving amide **24** (89 mg, 90%) as a clear oil. **^1^H NMR** (401 MHz, DMSO*-d*_6_) δ 8.16 (s, 1H), 7.55 (s, 1H), 7.43–7.24 (m, 12H), 6.07 (d, *J* = 6.7 Hz, 1H), 4.77 (s, 4H), 4.50 (dd, *J* = 6.8, 4.3 Hz, 1H), 4.06 (dd, *J* = 4.3, 1.8 Hz, 1H), 3.97–3.87 (m, 2H), 2.59–2.37* (m, 4H), 1.86–1.76 (m, 2H), 1.39–1.34 (m, 19H), 0.90 (s, 9H), 0.64 (s, 9H), 0.10 (s, 3H), 0.08 (s, 3H), -0.15 (s, 3H), -0.43 (s, 3H). **^13^C NMR** (101 MHz, DMSO*-d*_6_) δ 171.6, 167.0, 158.1, 155.7, 153.1, 151.0, 137.0, 128.9, 127.6, 123.6, 109.3, 102.2, 87.0, 84.4, 80.6, 78.3, 74.5, 74.4, 53.5, 48.3*, 33.3, 31.1, 28.3, 28.3, 27.8, 25.9, 25.6, 17.9, 17.6, -4.5, -4.5, -4.6, -5.4. **LRMS** (ESI) [M + H]^+^ m/z: 991.5.

***S*-(((2*S*,3*S*,4*R*,5*R*)-5-(4-Amino-5-(dibenzylcarbamoyl)-7*H*-pyrrolo[2,3-*d*]pyrimidin-7-yl)-3,4-dihydroxytetrahydrofuran-2-yl)methyl)-l-homocysteine (STM1189)**

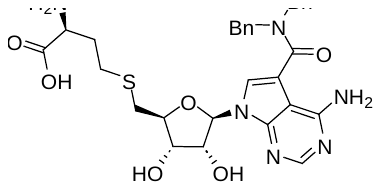
Compound **24** (87 mg, 0.088 mmol) was dissolved in 90% aqueous TFA (3.0 mL) and stirred at RT for 6 h. After complete deprotection, volatiles were removed under reduced pressure and the residue was purified by RP-FCC (10–70% of ACN in water, 0.1% FA), yielding compound **STM1189** (43 mg, 81%) as a clear solid. **^1^H NMR** (401 MHz, DMSO*-d*_6_) δ 8.15 (s, 1H), 7.46–7.23 (m, 13H), 6.05 (d, *J* = 5.2 Hz, 1H), 4.91–4.64 (m, 4H), 4.13 (t, *J* = 5.1 Hz, 1H), 3.89 (q, *J* = 6.1 Hz, 1H), 3.79–3.71 (m, 1H), 3.32–3.24 (m, 1H), 2.61–2.51 (m, 3H), 2.29 (dd, *J* = 13.8, 6.6 Hz, 1H), 2.02–1.89 (m, 1H), 1.85–1.72 (m, 1H). **^13^C NMR** (101 MHz, DMSO*-d*_6_) δ 169.9, 167.2, 158.1, 153.1, 150.8, 137.2, 129.0, 127.6, 123.4, 109.1, 101.9, 87.2, 83.0, 74.0, 72.8, 53.2, 49.3*, 34.0, 31.5, 28.3. **HRMS** (ESI) m/z: [M + H]^+^ (C_30_H_35_N_6_O_6_S) calculated 607.2333, found 607.2335.

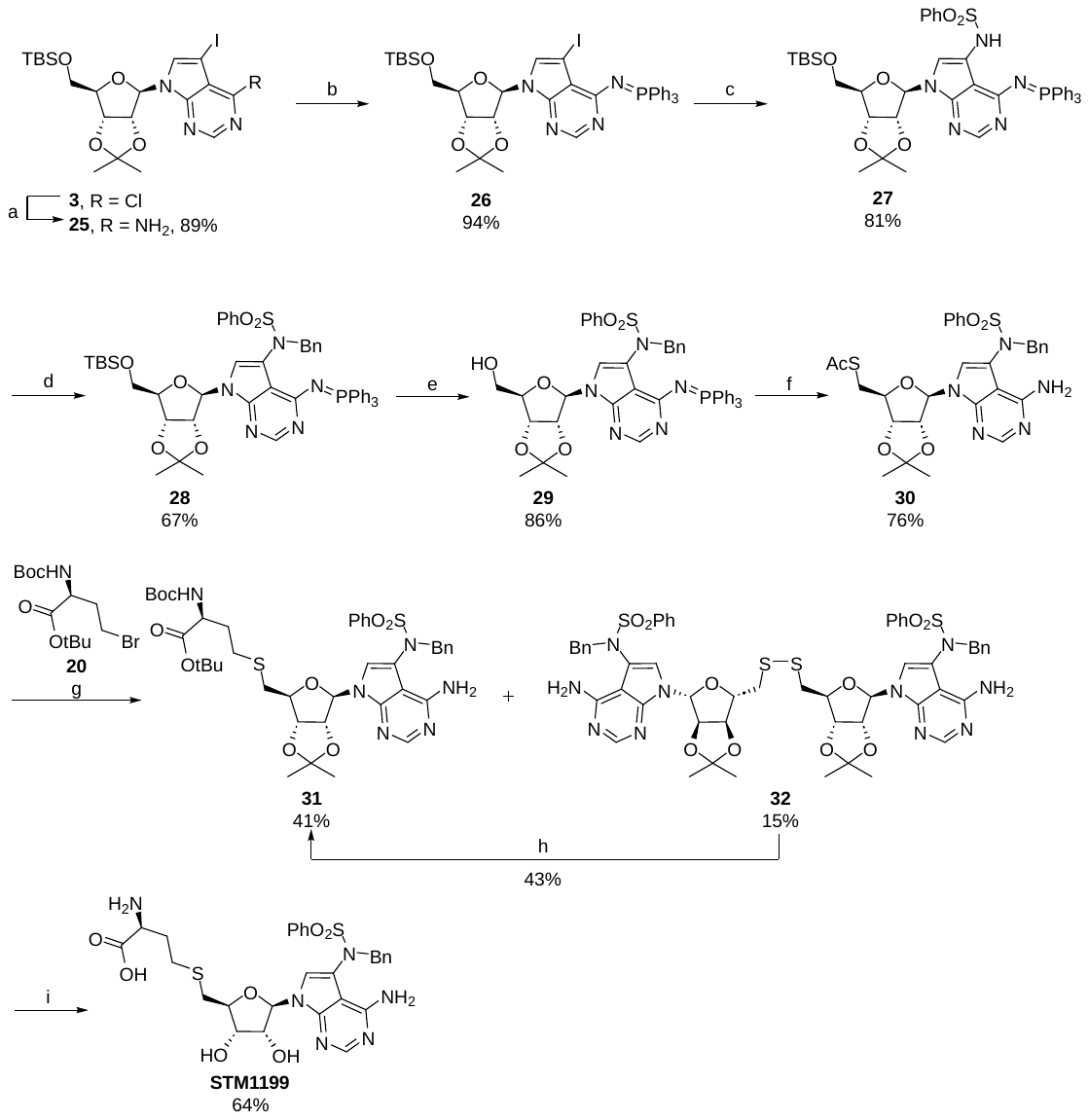

**SI Scheme 4** – Reagents and conditions: (a) 4 M NH_3_ in EtOH, RT, 20 h; (b) PPh_3_, DIAD, THF, RT, overnight; (c) PhSO_2_NH_2_, CuI, DMDACH, K_3_PO_4_, 1,4-dioxane, 110 °C, 6 h; (d) BnBr, Cs_2_CO_3_, DMF, RT, 3 h; (e) TBAF (1 M in THF), THF, RT, 1h; (f) (i) AcSH, PPh_3_, DIAD, THF, 0 °C to RT, 90 min, (ii) AcOH, H_2_O, THF, 65 °C, 5 h; (g) **20**, DBU, PPh_3_, MeOH, RT, 20 h; (h) **20**, NaBH_4_, DMF/EtOH (2:3) mixture, RT, 10 h; (i) 90% TFA (aq.), RT, 9 h.

**7-((3a*R*,4*R*,6*R*,6a*R*)-6-(((*tert*-Butyldimethylsilyl)oxy)methyl)-2,2-dimethyltetrahydro**

**furo[3,4-*d*][1,3]dioxol-4-yl)-5-iodo-7*H*-pyrrolo[2,3-*d*]pyrimidin-4-amine (25)**

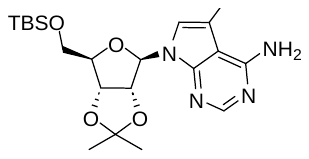
The modified nucleoside **3** (22.0 g, 38.9 mmol), prepared according to a reported procedure,[7] was transferred to a pressure tube and dissolved in 4 M solution of NH_3_ in EtOH (150 mL). The reaction mixture was then heated to 100 °C and stirred for 12 h. After removing volatiles in vacuo, the residue was dissolved in DCM and washed with water. The organic layer was dried over Na_2_SO_4_, concentrated under reduced pressure, and subjected to column chromatography (30–70% of EtOAc in toluene) to obtain compound **25** (18.95 g, 89 %) as a brownish foam. NMR characteristics were consistent with the published data.[8]

***N*-(7-((3a*R*,4*R*,6*R*,6a*R*)-6-(((*tert*-Butyldimethylsilyl)oxy)methyl)-2,2-dimethyltetrahydrofuro[3,4-*d*][1,3]dioxol-4-yl)-5-iodo-7*H*-pyrrolo[2,3-*d*]pyrimidin-4-yl)-1,1,1-triphenyl-λ^5^-phosphanimine (26)**

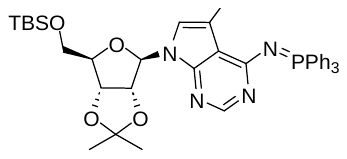
Starting material **25** (1.00 g, 1.83 mmol) and PPh_3_ (672 mg, 2.56 mmol, 1.4 eq) were dissolved in anhydrous THF (18.3 mL) and DIAD (467 µL, 2.38 mmol, 1.3 eq) was added at RT. The reaction mixture was stirred at RT overnight. Volatiles were removed under reduced pressure, the residue was adsorbed onto silica and purified by FCC (5–35% of EtOAc in cyclohexane) to afford compound **26** (1.38 g, 1.71 mmol, 94%) as a white foam. **^1^H NMR** (400 MHz, DMSO*-d*_6_) δ 7.97–7.88 (m, 6H), 7.87 (s, 1H), 7.67–7.60 (m, 3H), 7.59–7.52 (m, 6H), 7.52 (s, 1H), 6.18 (d, *J* = 2.8 Hz, 1H), 5.13 (dd, *J* = 6.3, 2.8 Hz, 1H), 4.88 (dd, *J* = 6.3, 3.1 Hz, 1H), 4.15–4.08 (m, 1H), 3.73 (dd, *J* = 11.1, 4.7 Hz, 1H), 3.67 (dd, *J* = 11.1, 5.0 Hz, 1H), 1.51 (s, 3H), 1.30 (s, 3H), 0.86 (s, 9H), 0.02 (s, 6H). **^13^C NMR** (101 MHz, DMSO*-d*_6_) δ 162.0 (d, *J* = 6.9 Hz), 151.3, 149.7 (d, *J* = 4.9 Hz), 133.1 (d, *J* = 10.0 Hz), 132.4, 128.91 (d, *J* = 99.6 Hz), 128.88 (d, *J* = 12.0 Hz), 126.8, 113.2, 110.4 (d, *J* = 23.7 Hz), 88.9, 85.6, 84.0, 80.9, 63.4, 55.2, 27.3, 26.0, 25.4, 18.2, -5.2, -5.2. **LRMS** (ESI) [M + H]^+^ m/z: 807.1.

***N*-(7-((3a*R*,4*R*,6*R*,6a*R*)-6-(((*tert*-Butyldimethylsilyl)oxy)methyl)-2,2-dimethyltetrahydrofuro[3,4-*d*][1,3]dioxol-4-yl)-4-((triphenyl-λ^5^-phosphaneylidene)amino)-7*H*-pyrrolo[2,3-*d*]pyrimidin-5-yl)benzenesulfonamide (27)**

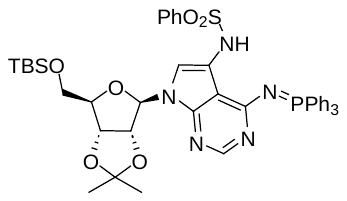
CuI (12 mg, 0.062 mmol, 1.0 eq) was placed in a screw-cap vial equipped with an argon balloon. Deoxygenated 1,4-dioxane (0.50 mL) and *trans*-*N,N*-dimethyl-1,2-diaminocyclohexane (20 µL, 0.12 mmol, 2.0 eq) were added. The resulting mixture was stirred at RT for 15 min, during which the suspension became clear and turned blue-green. In a separate vial, starting material **26** (50 mg, 0.062 mmol, 1.0 eq), PhSO_2_NH_2_ (39 mg, 0.25 mmol, 4.0 eq), and ground K_3_PO_4_ (27 mg, 0.12 mmol, 2.0 eq) were combined under argon. The copper complex solution was added, and the reaction mixture was stirred at 110 °C for 6 h in an aluminum heating block. After completion, volatiles were removed under reduced pressure, the residue was diluted with MeOH, adsorbed onto silica, and purified by RP-FCC (30–100% of ACN in water), furnishing compound **27** (42 mg, 81%). **^1^H NMR** (400 MHz, CDCl_3_) δ 8.86 (s, 1H), 7.93 (s, 1H), 7.70–7.56 (m, 10H), 7.53–7.41 (m, 7H), 7.39–7.30 (m, 1H), 7.17–7.08 (m, 2H), 7.05 (s, 1H), 6.20 (d, *J* = 3.4 Hz, 1H), 5.10 (dd, *J* = 6.5, 3.3 Hz, 1H), 4.91 (dd, *J* = 6.5, 3.2 Hz, 1H), 4.23–4.15 (m, 1H), 3.81–3.69 (m, 2H), 1.59 (s, 3H), 1.36 (s, 3H), 0.90 (s, 9H), 0.05 (s, 3H), 0.04 (s, 3H). **^13^C NMR** (101 MHz, CDCl_3_) δ 162.0*, 152.8, 148.4, 139.6, 133.1 (d, *J* = 10.1 Hz), 132.5 (d, *J* = 2.8 Hz), 132.4, 128.9 (d, *J* = 12.4 Hz), 128.8, 128.7 (d, *J* = 100.8 Hz), 127.2, 116.5, 114.3, 110.1, 105.0*, 89.6, 85.4, 84.2, 81.3, 63.5, 18.6, -5.2. **LRMS** (ESI) [M + H]^+^ m/z: 836.1.

***N*-Benzyl-*N*-(7-((3a*R*,4*R*,6*R*,6a*R*)-6-(((*tert*-butyldimethylsilyl)oxy)methyl)-2,2-dimethyltetrahydrofuro[3,4-*d*][1,3]dioxol-4-yl)-4-((triphenyl-λ^5^-phosphaneylidene)amino)-7*H*-pyrrolo[2,3-*d*]pyrimidin-5-yl)benzenesulfonamide (28)**

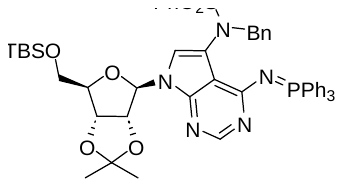
Starting material **27** (168 mg, 0.201 mmol) was dissolved in anhydrous DMF (4.0 mL). Cs_2_CO_3_ (131 mg, 0.40 mmol, 2.0 eq) and benzyl bromide (48 µL, 0.40 mmol, 2.0 eq) were added and the reaction mixture was stirred at RT under argon for 3 h. After completion, the mixture was concentrated, diluted with MeOH, adsorbed onto silica, and purified by RP-FCC (30–100% of ACN in water), giving compound **28** (124 mg, 67%). **^1^H NMR** (400 MHz, CDCl_3_) δ 7.94 (s, 1H), 7.81–7.74 (m, 2H), 7.66–7.52 (m, 10H), 7.49–7.40 (m, 6H), 7.27–7.17 (m, 4H), 7.17–7.10 (m, 1H), 7.08–7.01 (m, 2H), 7.00 (s, 1H), 6.24 (d, *J* = 2.8 Hz, 1H), 5.55–5.31 (m, 2H), 5.06–4.94 (m, 1H), 4.88 (dd, *J* = 6.4, 3.2 Hz, 1H), 4.23–4.05 (m, 1H), 3.79–3.65 (m, 2H), 1.60 (s, 3H), 1.36 (s, 3H), 0.89 (s, 9H), 0.01 (s, 3H), -0.01 (s, 3H). **^13^C NMR** (101 MHz, CDCl_3_) δ 161.7 (d, *J* = 7.3 Hz), 151.6, 150.0 (d, *J* = 5.3 Hz), 140.9, 137.8, 133.3 (d, *J* = 9.8 Hz), 132.0 (d, *J* = 2.8 Hz), 131.9, 129.4 (d, *J* = 100.2 Hz), 128.7, 128.47 (d, *J* = 12.1 Hz), 128.46, 128.4, 127.8, 127.3, 121.9, 115.0, 114.3, 107.4, 89.6, 85.7, 84.6, 81.3, 63.4, 55.5, 27.4, 26.1, 25.6, 18.5, -5.2, -5.3. **LRMS** (ESI) [M + H]^+^ m/z: 926.2.

***N*-Benzyl-*N*-(7-((3a*R*,4*R*,6*R*,6a*R*)-6-(hydroxymethyl)-2,2-dimethyltetrahydrofuro[3,4-*d*][1,3]dioxol-4-yl)-4-((triphenyl-λ^5^-phosphaneylidene)amino)-7*H*-pyrrolo[2,3-*d*]pyrimidin-5-yl)benzenesulfonamide (29)**

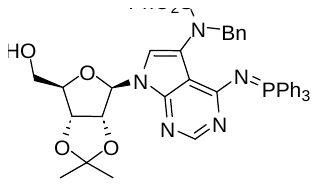
Intermediate **28** (124 mg, 0.134 mmol) was dissolved in THF (3.2 mL, dried over NaOH) and 1 M TBAF solution (235 µL, 0.235 mmol, 1.8 eq) was added. The reaction mixture was stirred at RT for 1 h until complete consumption of the starting material. The mixture was adsorbed onto silica and purified by RP-FCC (20–100% of ACN in water), furnishing compound **29** (93 mg, 86%) as a white solid. **^1^H NMR** (401 MHz, CDCl_3_) δ 7.85 (s, 1H), 7.83–7.76 (m, 2H), 7.68–7.54 (m, 9H), 7.50–7.41 (m, 6H), 7.32–7.29 (m, 1H), 7.27–7.21 (m, 3H), 7.21–7.14 (m, 1H), 7.11–7.04 (m, 2H), 6.92 (s, 1H), 6.80 (bs, 1H), 5.62 (d, *J* = 4.7 Hz, 1H), 5.50–5.34 (m, 2H), 5.20 (t, *J* = 5.3 Hz, 1H), 5.06 (dd, *J* = 6.0, 1.3 Hz, 1H), 4.45–4.39 (m, 1H), 3.91 (dd, *J* = 12.6, 1.4 Hz, 1H), 3.73 (dd, *J* = 12.6, 1.7 Hz, 1H), 1.60 (s, 3H), 1.34 (s, 3H). **^13^C NMR** (101 MHz, CDCl_3_) δ 162.2 (d, *J* = 7.8 Hz), 150.6, 147.7, 140.7, 137.6, 133.2 (d, *J* = 9.9 Hz), 132.0 (d, *J* = 2.7 Hz), 131.9, 129.3, 128.43, 128.42 (d, *J* = 12.3 Hz), 128.41, 128.3, 127.7, 127.3, 124.7, 113.8, 113.6, 109.1, 96.1, 85.5, 82.9, 81.4, 63.5, 55.5, 27.7, 25.3. **LRMS** (ESI) [M + H]^+^ m/z: 812.4.

***S*-(((3a*S*,4*S*,6*R*,6a*R*)-6-(4-Amino-5-(*N*-benzylphenylsulfonamido)-7*H*-pyrrolo[2,3-*d*]pyrimidin-7-yl)-2,2-dimethyltetrahydrofuro[3,4-*d*][1,3]dioxol-4-yl)methyl) ethanethioate (30)**

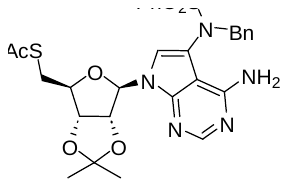
Compound **30** was prepared analogously to the synthesis of **19**, starting from compound **29** (93 mg, 0.115 mmol). Purification by RP-FCC (30–100% of ACN in water) provided compound **107** (60 mg, 86%). **^1^H NMR** (400 MHz, CDCl_3_) δ 8.23 (s, 1H), 7.80–7.73 (m, 2H), 7.71–7.62 (m, 1H), 7.61–7.51 (m, 2H), 7.24–7.15 (m, 5H), 6.31 (s, 1H), 5.96 (d, *J* = 2.0 Hz, 1H), 5.42 (s, 2H), 5.17–5.11 (m, 1H), 5.03 (bs, 1H), 4.71 (dd, *J* = 6.3, 3.2 Hz, 1H), 4.38 (bs, 1H), 4.19 (td, *J* = 7.0, 3.2 Hz, 1H), 3.06–2.91 (m, 2H), 2.35 (s, 3H), 1.55 (s, 3H), 1.34 (s, 3H). **^13^C NMR** (101 MHz, CDCl_3_) δ 194.7, 157.1, 153.0, 148.8, 135.4, 133.5, 129.2, 129.0, 128.8, 128.3, 128.3, 121.5, 115.3, 114.5, 102.5, 91.0, 85.7, 84.8, 83.5, 57.3, 31.3, 30.8, 27.2, 25.5. **LRMS** (ESI) [M + H]^+^ m/z: 610.1.

***tert*-Butyl *S*-(((3a*S*,4*S*,6*R*,6a*R*)-6-(4-amino-5-(*N*-benzylphenylsulfonamido)-7*H*-pyrrolo[2,3-*d*]pyrimidin-7-yl)-2,2-dimethyltetrahydrofuro[3,4-*d*][1,3]dioxol-4-yl)methyl)-*N*-(*tert*-butoxycarbonyl)-l-homocysteinate (31)**

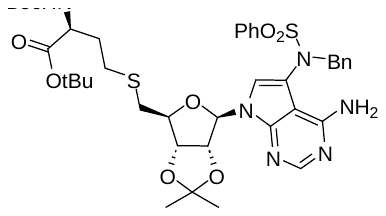
Starting material **30** (58 mg, 0.095 mmol) was dissolved in anhydrous MeOH (2.0 mL, previously deoxygenated by bubbling with argon). PPh_3_ (3 mg, 0.011 mmol, 0.12 eq) and bromide **20** (39 mg, 0.114 mmol, 1.2 eq) were added, followed by DBU (31 µL, 0.21 mmol, 2.2 eq). The reaction mixture was stirred at RT under argon and an additional portion of bromide **20** (39 mg, 0.114 mmol, 1.2 eq) was added after 7 h. Stirring was continued overnight. The crude mixture, containing desired product **31**, disulfide **32**, and partially deprotected side product **31-DeBoc,** was concentrated under reduced pressure, and the residue was subjected to RP-FCC (20–100% of ACN in water). Since separation of the desired product from the disulfide was not achieved, impure fractions were concentrated and further purified by FCC (5–50% of EtOAc/EtOH (4:1) in DCM), providing compound **31** (32 mg, 0.039 mmol, 41%), disulfide **32** (8 mg, 0.007 mmol, 15%), and **31-DeBoc** (8 mg, 0.011 mmol, 12%).

**Use of side product 32 for the preparation of 31.** Disulfide **32** (8 mg, 0.007 mmol) and bromide **20** (5 mg, 0.015 mmol, 2.1 eq) were dissolved in a mixture of DMF (0.40 mL) and EtOH (0.60 mL). The solution was deoxygenated by repeated evacuation–refilling with argon, and NaBH_4_ (1.3 mg, 0.035 mmol, 5.0 eq) was added. The reaction mixture was stirred at RT under argon for 5 h then an additional portion of bromide **20** (5 mg, 0.015 mmol, 2.1 eq) was added and stirring was continued overnight. After completion, volatiles were removed under reduced pressure and the residue was purified by RP-FCC (30–100% of ACN in water, 0.1% FA), furnishing compound **31** (5 mg, 43%). **^1^H NMR** (401 MHz, CDCl_3_) δ 8.21 (s, 1H), 7.79–7.72 (m, 2H), 7.71–7.63 (m, 1H), 7.59–7.51 (m, 2H), 7.25–7.14 (m, 5H), 6.30 (s, 1H), 5.95 (s, 1H), 5.42 (s, 2H), 5.19–4.95 (m, 3H), 4.86–4.79 (m, 1H), 4.48–4.19 (m, 3H), 2.70–2.57 (m, 2H), 2.57–2.44 (m, 2H), 2.07–1.94 (m, 1H), 1.87–1.75 (m, 1H), 1.57 (s, 3H), 1.45 (s, 9H), 1.44 (s, 9H), 1.36 (s, 3H). **^13^C NMR** (101 MHz, CDCl_3_) δ 171.4, 157.0, 152.8, 148.7, 137.3, 135.4, 133.6, 129.2, 129.0, 128.8, 128.4, 128.2, 121.7, 115.3, 114.6, 102.5, 91.0, 85.6, 84.7, 83.5, 82.4, 57.4, 53.4, 34.5, 33.2, 28.5, 28.3*, 28.1, 27.2, 25.5. **LRMS** (ESI) [M + H]^+^ m/z: 825.3.

***S*-(((2*S*,3*S*,4*R*,5*R*)-5-(4-Amino-5-(*N*-benzylphenylsulfonamido)-7*H*-pyrrolo[2,3-*d*]pyrimidin-7-yl)-3,4-dihydroxytetrahydrofuran-2-yl)methyl)-l-homocysteine (STM1199)**

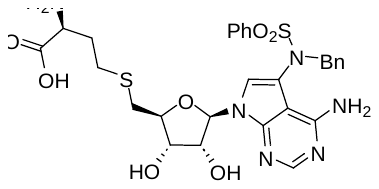
Both isolated intermediates, **31** (39 mg, 0.047 mmol) and **31-DeBoc** (9 mg, 0.012 mmol) were dissolved in 90% aqueous TFA (1.5 mL) and the resulting mixture was stirred at RT for 9 h. Volatiles were removed under reduced pressure, and the residue was co-evaporated with EtOH. Purification by RP-FCC (15–50% of ACN in water, 0.1% FA) gave compound **STM1199** (24 mg, 64%) as a white solid. **^1^H NMR** (600 MHz, DMSO, 70 °C) δ 8.04 (s, 1H), 7.82–7.78 (m, 1H), 7.75–7.71 (m, 2H), 7.68–7.63 (m, 2H), 7.27–7.16 (m, 5H), 6.81 (s, 1H), 6.24 (s, 2H), 5.99 (d, *J* = 4.8 Hz, 1H), 4.11–4.08 (m, 1H), 3.97–3.92 (m, 1H), 3.89–3.85 (m, 1H), 3.31–3.26 (m, 1H), 2.76 (dd, *J* = 13.7, 5.8 Hz, 1H), 2.68–2.62 (m, 2H), 2.60 (dd, *J* = 13.6, 6.1 Hz, 1H), 2.05–1.97 (m, 1H), 1.85–1.79 (m, 1H). **^13^C NMR** (151 MHz, DMSO*-d*_6_, 70 °C) δ 170.0, 156.8, 152.0, 148.6, 136.4, 135.7, 133.5, 129.1, 128.5, 128.2, 127.8, 127.6, 120.2, 114.8, 101.0, 87.1, 82.6, 73.9*, 72.4, 56.3, 53.2, 34.0, 32.0, 28.6. **HRMS** (ESI) m/z: [M + H]^+^ (C_28_H_33_N_6_O_7_S_2_) calculated 629.1847, found 629.1850.

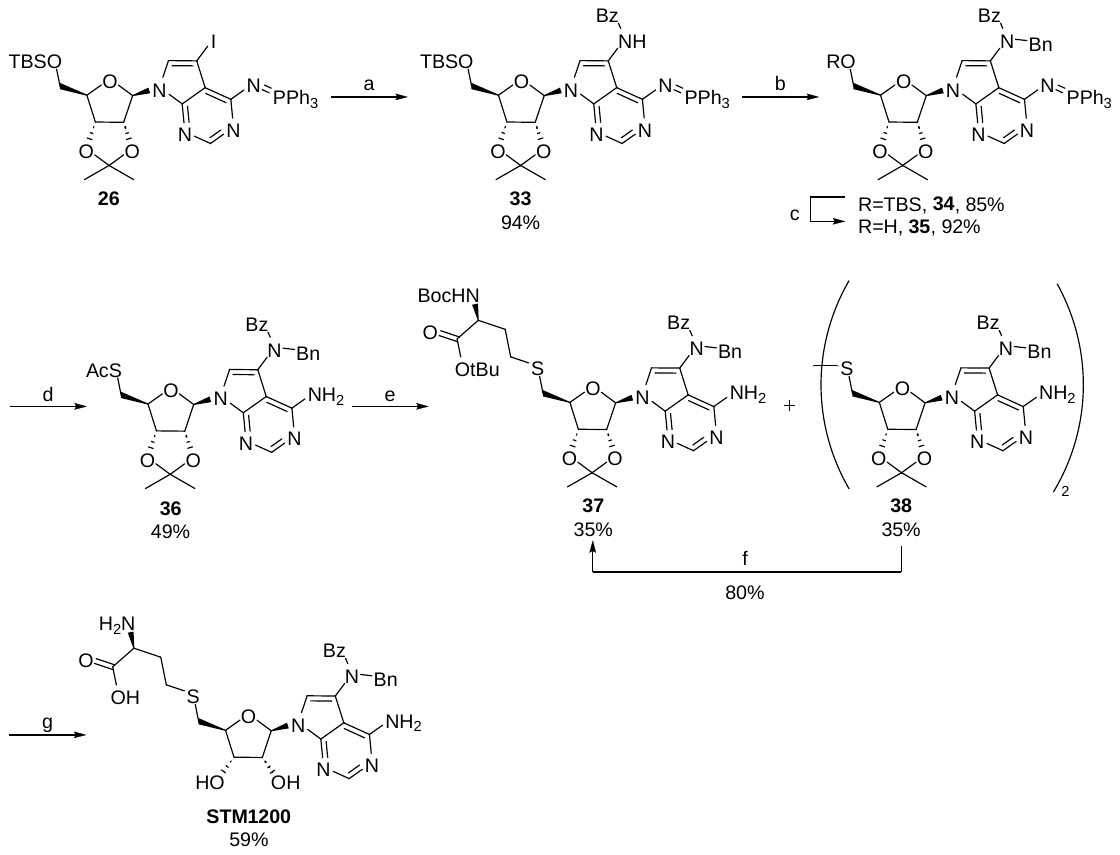

**SI Scheme 5** – Reagents and conditions: (a) BzNH_2_, CuI, DMDACH, K_3_PO_4_, 1,4-dioxane, 110 °C, 1h; (b) BnBr, NaH, DMF, RT, 90 min, (c) TBAF (1 M in THF), THF, RT, 1h; (d) (i) AcSH, PPh_3_, DIAD, THF, 0 °C to RT, 90 min, (ii) AcOH, H_2_O, THF, 65 °C, 5 h; (e) **20**, DBU, PPh_3_, MeOH, RT, 20 h; (f) **99**, NaBH_4_, DMF/EtOH (2:3) mixture, RT, 10 h; (g) 90% TFA (aq.), RT, 9 h.

***N*-(7-((3a*R*,4*R*,6*R*,6a*R*)-6-(((*tert*-Butyldimethylsilyl)oxy)methyl)-2,2-dimethyltetrahydrofuro[3,4-*d*][1,3]dioxol-4-yl)-4-((triphenyl-λ^5^-phosphaneylidene)amino)-7*H*-pyrrolo[2,3-*d*]pyrimidin-5-yl)benzamide (33)**

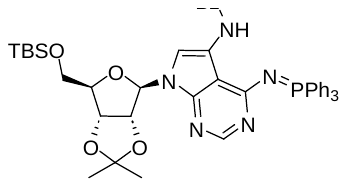
CuI (47 mg, 0.25 mmol, 1.0 eq) was placed into a flask equipped with an argon-filled balloon and deoxygenated dioxane (2.0 mL) was added, followed by *trans*-*N,N*-dimethyl-1,2-diaminocyclohexane (0.078 mL, 0.50 mmol, 2.0 eq). The resulting suspension was stirred at RT for 10 min until complete dissolution of CuI. Benzamide (120 mg, 0.99 mmol, 4.0 eq), compound **26** (200 mg, 0.25 mmol), and ground K_3_PO_4_ (105 mg, 0.50 mmol, 2.0 eq) were added and the reaction mixture was stirred at 110 °C for 1 h. The reaction mixture was diluted with MeOH and adsorbed onto silica. The crude product was purified by RP-FCC (30–100% of ACN in water) to afford compound **33** (186 mg, 94%). **^1^H NMR** (400 MHz, CDCl_3_) δ 10.35 (s, 1H), 8.03 (s, 1H), 7.85–7.75 (m, 9H), 7.61–7.54 (m, 3H), 7.51–7.40 (m, 7H), 7.33–7.27 (m, 2H), 6.32 (d, *J* = 3.2 Hz, 1H), 5.27 (dd, *J* = 6.6, 3.2 Hz, 1H), 4.93 (dd, *J* = 6.6, 3.5 Hz, 1H), 4.19 (td, *J* = 5.6, 3.6 Hz, 1H), 3.76 (d, *J* = 5.7 Hz, 2H), 1.60 (s, 3H), 1.37 (s, 3H), 0.87 (s, 9H), 0.02 (s, 6H). **^13^C NMR** (101 MHz, CDCl_3_) δ 164.2, 162.7 (d, *J* = 7.3 Hz), 152.3, 148.3 (d, *J* = 5.2 Hz), 134.7, 133.3 (d, *J* = 9.9 Hz), 132.3 (d, *J* = 2.8 Hz), 131.2, 129.6, 129.1 (d, *J* = 100.3 Hz), 128.80 (d, *J* = 12.2 Hz), 128.75, 127.2, 117.9, 114.3, 109.7, 104.01 (d, *J* = 22.7 Hz), 89.4, 85.5, 84.0, 81.7, 63.6, 27.4, 26.1, 25.6, 18.6, -5.2. **LRMS** (ESI) [M + H]^+^ m/z: 800.2.

***N*-Benzyl-*N*-(7-((3a*R*,4*R*,6*R*,6a*R*)-6-(((*tert*-butyldimethylsilyl)oxy)methyl)-2,2-dimethyltetrahydrofuro[3,4-*d*][1,3]dioxol-4-yl)-4-((triphenyl-λ^5^-phosphaneylidene)amino)-7*H*-pyrrolo[2,3-*d*]pyrimidin-5-yl)benzamide (34)**

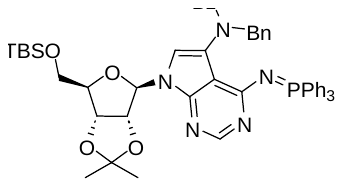
Starting material **33** (39 mg, 0.049 mmol) was dissolved in anhydrous DMF (1.0 mL) and NaH (3.9 mg, 0.098 mmol, 2.0 eq) was added in one portion. After cessation of gas evolution, benzyl bromide (17 µL, 0.146 mmol, 3.0 eq) was added and the reaction mixture was stirred at RT for 90 min. After completion, volatiles were removed under reduced pressure, and the residue was purified by RP-FCC (30–100% of ACN in water), to afford compound **34** (37 mg, 85%) as a clear solid. **^1^H NMR** (401 MHz, CDCl_3_) δ 7.97 (s, 1H), 7.91–7.78 (m, 6H), 7.60–7.37 (m, 11H), 7.34–7.29 (m, 2H), 7.23–7.16 (m, 3H), 7.14–7.05 (m, 1H), 7.00–6.91 (m, 2H), 6.36–6.18 (m, 1H), 6.07 (s, 1H), 5.96–5.80 (m, 1H), 4.89–4.40 (m, 3H), 4.03 (s, 1H), 3.65–3.48 (m, 2H), 1.59 (s, 2H), 1.53 (s, 3H), 0.85 (s, 9H), -0.04 (s, 3H), -0.08 (s, 3H). The ^13^C NMR spectrum of compound **34** was not recorded. **LRMS** (ESI) [M + H]^+^ m/z: 890.2.

***N*-Benzyl-*N*-(7-((3a*R*,4*R*,6*R*,6a*R*)-6-(hydroxymethyl)-2,2-dimethyltetrahydrofuro[3,4-*d*][1,3]dioxol-4-yl)-4-((triphenyl-λ^5^-phosphaneylidene)amino)-7*H*-pyrrolo[2,3-*d*]pyrimidin-5-yl)benzamide (35)**

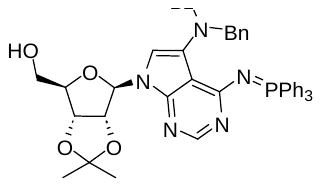
Starting material **34** (125 mg, 0.14 mmol) was dissolved in THF (3.4 mL, dried over NaOH) and 1 M TBAF solution (211 µL, 0.21 mmol, 1.5 eq) was added. The reaction mixture was stirred at RT for 1 h until complete consumption of the starting material. The mixture was adsorbed onto silica and purified by RP-FCC (20–100% of ACN in water), yielding compound **35** (100 mg, 0.13 mmol, 92%) as a white solid. **^1^H NMR** (401 MHz, CDCl_3_) δ 7.89–7.78 (m, 7H), 7.59–7.53 (m, 3H), 7.52–7.40 (m, 8H), 7.37–7.33 (m, 2H), 7.25–7.18 (m, 3H), 7.16–7.04 (m, 1H), 7.04–6.93 (m, 2H), 6.70 (s, 1H), 6.24 (d, *J* = 18.2 Hz, 1H), 5.92 (dd, *J* = 33.3, 13.8 Hz, 1H), 5.41 (s, 1H), 4.97 (s, 2H), 4.59 (dd, *J* = 34.6, 13.9 Hz, 1H), 4.34 (s, 1H), 3.85 (d, *J* = 12.4 Hz, 1H), 3.67 (d, *J* = 12.4 Hz, 1H), 1.52 (s, 3H), 1.29 (s, 3H). **^13^C NMR** (101 MHz, CDCl_3_) δ 171.6*, 162.7 (d, *J* = 7.1 Hz), 151.5, 138.1*, 137.0, 133.4 (d, *J* = 9.9 Hz), 132.2 (d, *J* = 2.7 Hz), 129.5, 129.3, 129.1, 128.7 (d, *J* = 12.3 Hz), 128.5, 128.4, 127.4, 127.2, 121.3, 113.6, 109.3, 96.0, 85.8, 83.2, 81.4, 63.5, 54.1, 27.7, 25.4. **LRMS** (ESI) [M + H]^+^ m/z: 776.1.

***S*-(((3a*S*,4*S*,6*R*,6a*R*)-6-(4-Amino-5-(*N*-benzylbenzamido)-7*H*-pyrrolo[2,3-*d*]pyrimidin-7-yl)-2,2-dimethyltetrahydrofuro[3,4-*d*][1,3]dioxol-4-yl)methyl) ethanethioate (36)**

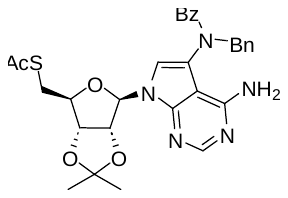
Starting material **35** (88 mg, 0.113 mmol) and PPh_3_ (66 mg, 0.25 mmol, 2.2 eq) were dissolved in anhydrous THF (1.1 mL) and the solution was cooled to 0 °C in an ice bath. DIAD (0.049 mL, 0.25 mmol, 2.2 eq) was added and the reaction mixture was stirred at 0 °C for 30 min. Thioacetic acid (0.018 mL, 0.25 mmol, 2.2 eq) was then added and stirring was continued for 1 h while allowing the temperature to rise to RT. Water (0.189 mL) and acetic acid (39 µL, 0.68 mmol, 6.0 eq) were added and the reaction mixture was stirred at 65 °C for 5 h to hydrolyze the formed iminophosphorane. Volatiles were removed under reduced pressure, and the residue was purified by RP-FCC (30–100% of ACN in water), yielding compound **36** (32 mg, 49%). **^1^H NMR** (401 MHz, CDCl_3_) δ 8.25 (s, 1H), 7.38–7.27 (m, 7H), 7.23–7.16 (m, 1H), 7.16–7.07 (m, 2H), 6.45 (s, 1H), 6.00 (s, 1H), 5.39–5.25 (m, 1H), 5.13 (s, 2H), 5.06 (d, *J* = 5.7 Hz, 1H), 4.79–4.74 (m, 1H), 4.68–4.63 (m, 1H), 4.15 (td, *J* = 6.9, 3.0 Hz, 1H), 2.84–2.78 (m, 2H), 2.36 (s, 3H), 1.53 (s, 3H), 1.32 (s, 3H). **^13^C NMR** (101 MHz, CDCl_3_) δ 194.7, 172.0*, 156.4, 153.1, 149.1, 136.7, 135.4, 130.1, 129.4, 128.9, 128.3, 128.0, 121.2, 118.9, 114.4, 101.1, 90.6, 85.7, 84.7, 83.4, 54.5, 31.1, 30.7, 27.1, 25.5. **LRMS** (ESI) [M + H]^+^ m/z: 574.3.

***tert*-Butyl *S*-(((3a*S*,4*S*,6*R*,6a*R*)-6-(4-amino-5-(*N*-benzylbenzamido)-7*H*-pyrrolo[2,3-*d*]pyrimidin-7-yl)-2,2-dimethyltetrahydrofuro[3,4-*d*][1,3]dioxol-4-yl)methyl)-*N*-(*tert*-butoxycarbonyl)-l-homocysteinate (37)**

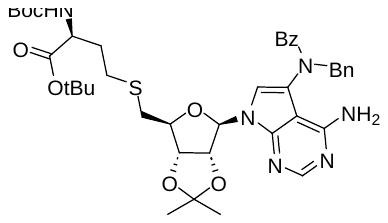
Starting material **36** (31 mg, 0.054 mmol) was dissolved in anhydrous MeOH (1.2 mL, previously deoxygenated by bubbling with argon). PPh_3_ (2 mg, 0.008 mmol, 0.14 eq) and bromide **20** (22 mg, 0.065 mmol, 1.2 eq) were added, followed by DBU (18 µL, 0.12 mmol, 2.2 eq). The reaction mixture was stirred under argon at RT for 20 h. The crude mixture, containing desired product, disulfide **38**, and partially deprotected side product **37-DeBoc**, was adsorbed onto silica and subjected to FCC (10–100% of EtOAc/EtOH (4:1) in DCM). Fractions were further purified by RP-FCC (30–100% of ACN in water, 0.1% FA), furnishing compound **37** (15 mg, 35%), disulfide **38** (10 mg, 35%), and **37-DeBoc** (5 mg, 13%).

**Use of 38 for the preparation of 37.** Disulfide **38** (10 mg, 0.009 mmol) and bromide **20** (6 mg, 0.018 mmol, 1.9 eq) were dissolved in a mixture of DMF (0.40 mL) and EtOH (0.60 mL). The solution was deoxygenated by repeated evacuation–refilling with argon, and NaBH_4_ (2 mg, 0.047 mmol, 5.0 eq) was added. The reaction mixture was stirred at RT under argon for 5 h then additional bromide **20** (3 mg, 0.009 mmol, 0.9 eq) was added and stirring was continued for another 5 h. After consumption of the starting material, volatiles were removed under reduced pressure and the residue was purified by RP-FCC (30–100% of ACN in water), providing compound **37** (12 mg, 80%). **100**: **^1^H NMR** (400 MHz, CDCl_3_) δ 8.22 (s, 1H), 7.40–7.28 (m, 7H), 7.24–7.08 (m, 3H), 6.45 (s, 1H), 5.98 (s, 1H), 5.41–5.20 (m, 1H), 5.17–5.05 (m, 3H), 5.04–4.93 (m, 1H), 4.78 (s, 1H), 4.27–4.15 (m, 2H), 2.57–2.26 (m, 4H), 2.00 (s, 1H), 1.82 (s, 1H), 1.55 (s, 3H), 1.46 (s, 9H), 1.44 (s, 9H), 1.33 (s, 3H). **^13^C NMR** (101 MHz, CDCl_3_) δ 171.4, 156.3, 155.5, 152.9, 149.0, 136.7, 135.4, 130.2, 129.4, 129.0, 128.3, 128.1, 121.3*, 114.5, 90.7, 85.7, 84.7, 83.5, 82.4, 80.0, 54.6*, 53.4, 34.3, 33.2, 28.5, 28.4, 28.1, 27.2, 25.5. **LRMS** (ESI) [M + H]^+^ m/z: 789.2.

***S*-(((2*S*,3*S*,4*R*,5*R*)-5-(4-Amino-5-(*N*-benzylbenzamido)-7*H*-pyrrolo[2,3-*d*]pyrimidin-7-yl)-3,4-dihydroxytetrahydrofuran-2-yl)methyl)-l-homocysteine (STM1200)**

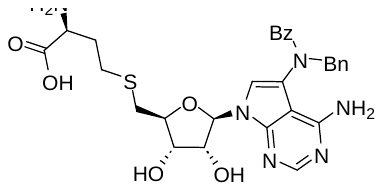
Both isolated intermediates, **37** (21 mg, 0.027 mmol) and **37-DeBoc** (5 mg, 0.007 mmol), were dissolved in 90% aqueous TFA (1.5 mL) and the resulting mixture was stirred at RT for 9 h. Volatiles were removed under reduced pressure, and the residue was co-evaporated with EtOH. Purification by RP-FCC (15–40% of ACN in water, 0.1% FA) furnished the deprotected product **STM1200** (12 mg, 59%) as a white solid. NMR analysis in DMSO-*d*_6_ indicated that the compound exists in solution as an approximately 1:1 mixture of rotamers. **^1^H NMR** (400 MHz, DMSO) δ 8.03 (s, 1H), 7.39–7.14 (m, 10H), 6.83 (s, 1H), 6.74 (s, 0.5H), 6.70 (s, 0.5H), 5.84 (d, *J* = 2.9 Hz, 1H), 5.59 (dd, *J* = 15.1, 5.4 Hz, 1H), 4.30 (d, *J* = 14.5 Hz, 1H), 4.01–3.97 (m, 0.5H), 3.86–3.74 (m, 2H), 3.73–3.67 (m, 0.5H), 3.31–3.24 (m, 1H), 2.75–2.61 (m, 1H), 2.61–2.30 (m, 3H), 2.05–1.90 (m, 1H), 1.87–1.73 (m, 1H). **^13^C NMR** (101 MHz, DMSO) δ 170.9, 169.9, 156.7, 152.4, 149.2, 149.2, 137.0, 136.5, 129.4, 129.4, 128.6, 128.5, 128.4, 127.9, 127.8, 127.4, 121.0, 120.9, 118.2, 118.1, 99.3, 99.0, 87.4, 87.2, 82.4, 82.2, 74.0, 73.7, 72.5, 72.4, 53.4, 53.3, 53.1, 33.7, 31.6, 28.4. **HRMS** (ESI) m/z: [M + H]^+^ (C_29_H_33_N_6_O_6_S) calculated 593.2177, found 593.2178.

### ^1^H and ^13^C NMR spectra of final compounds

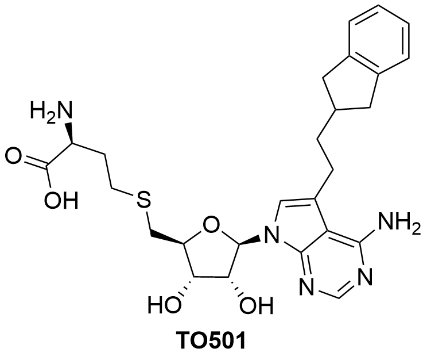

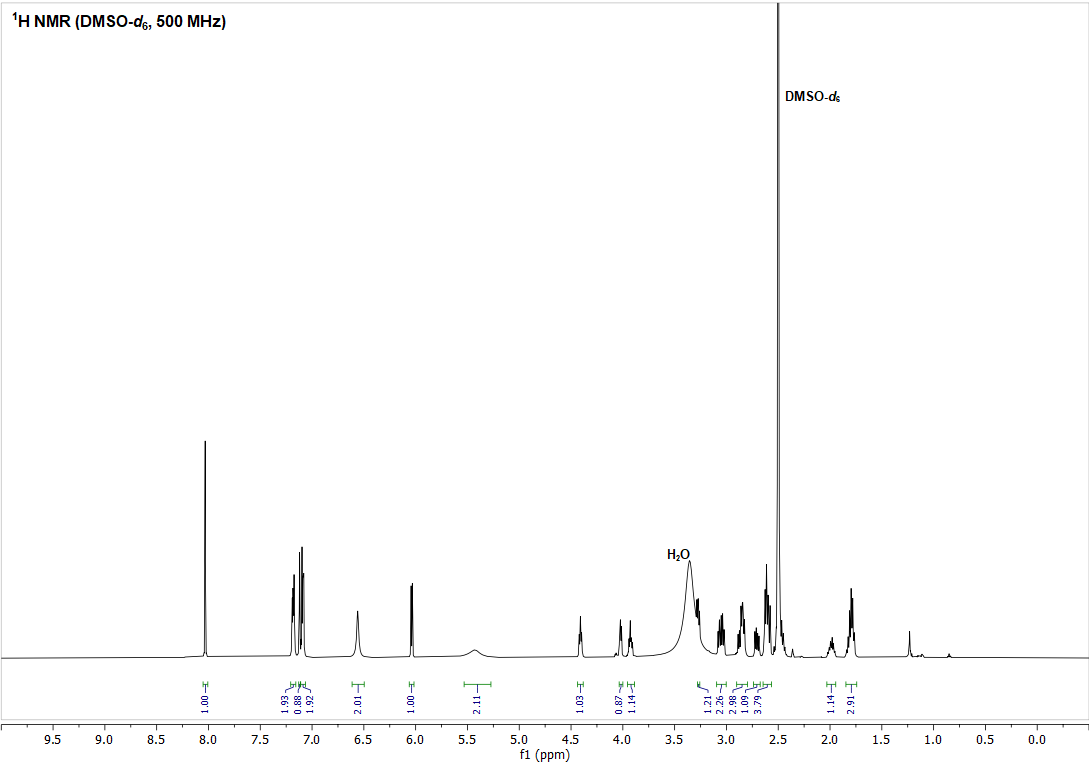

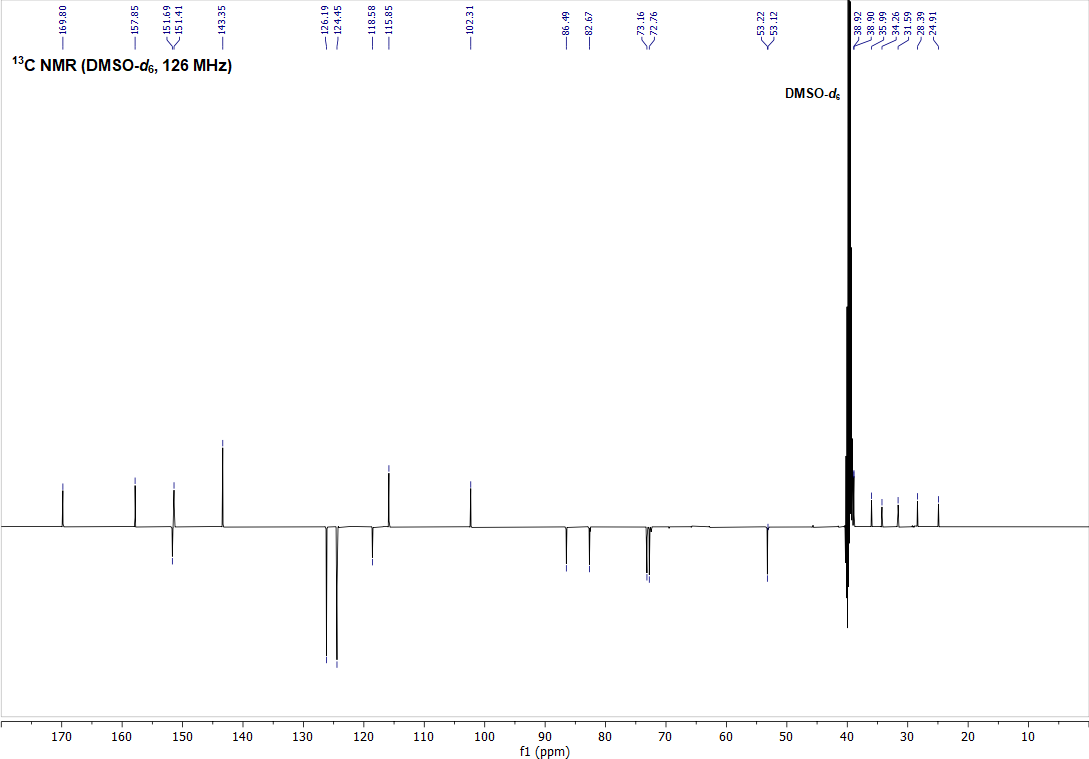

**SI Figure 1** – ^1^H (top) and ^13^C APT (bottom) NMR spectra of compound **TO501** measured in DMSO-*d*_6_.

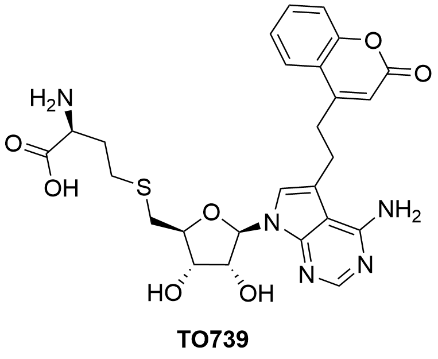

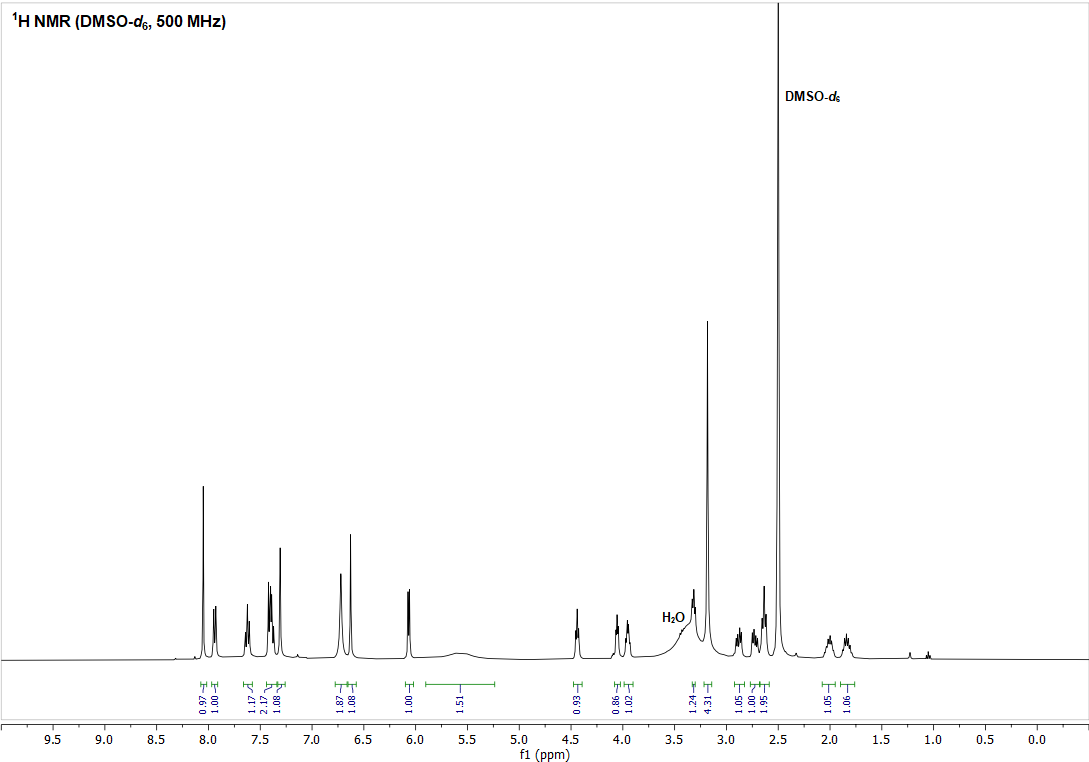

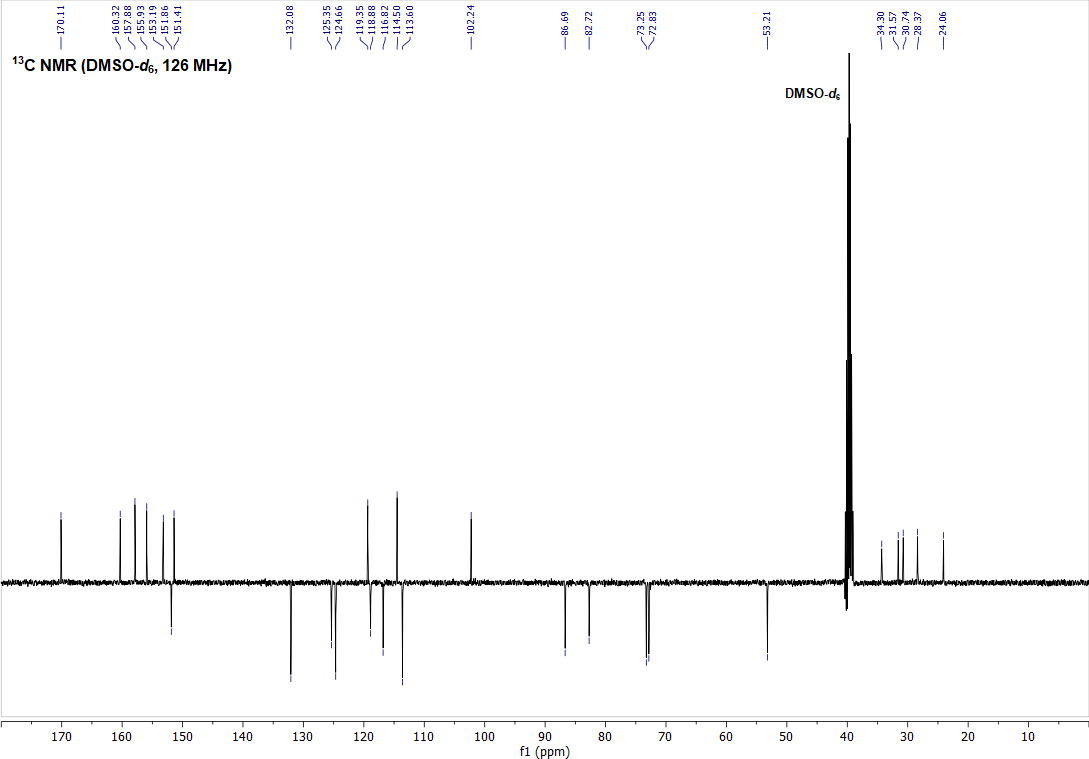

**SI Figure 2** – ^1^H (top) and ^13^C APT (bottom) NMR spectra of compound **TO739** measured in DMSO-*d*_6_.

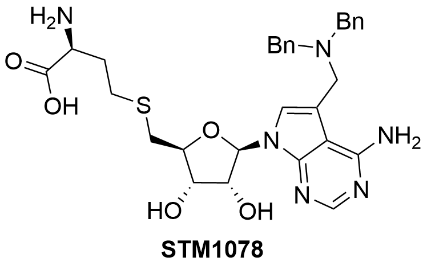

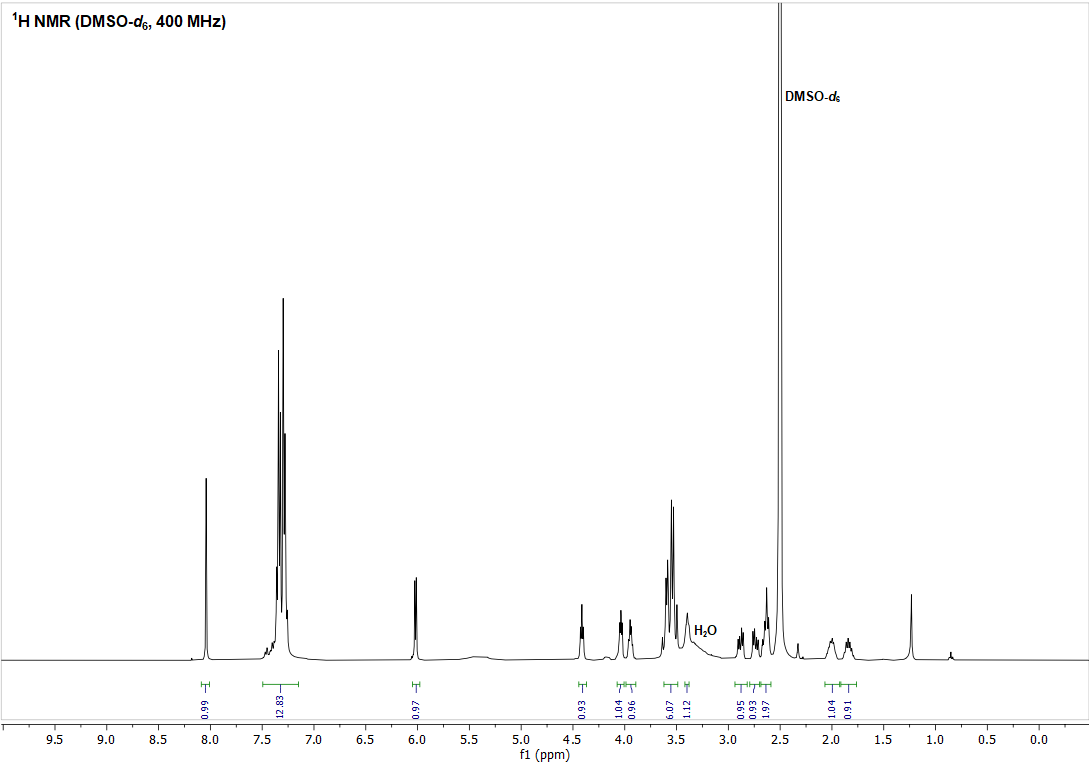

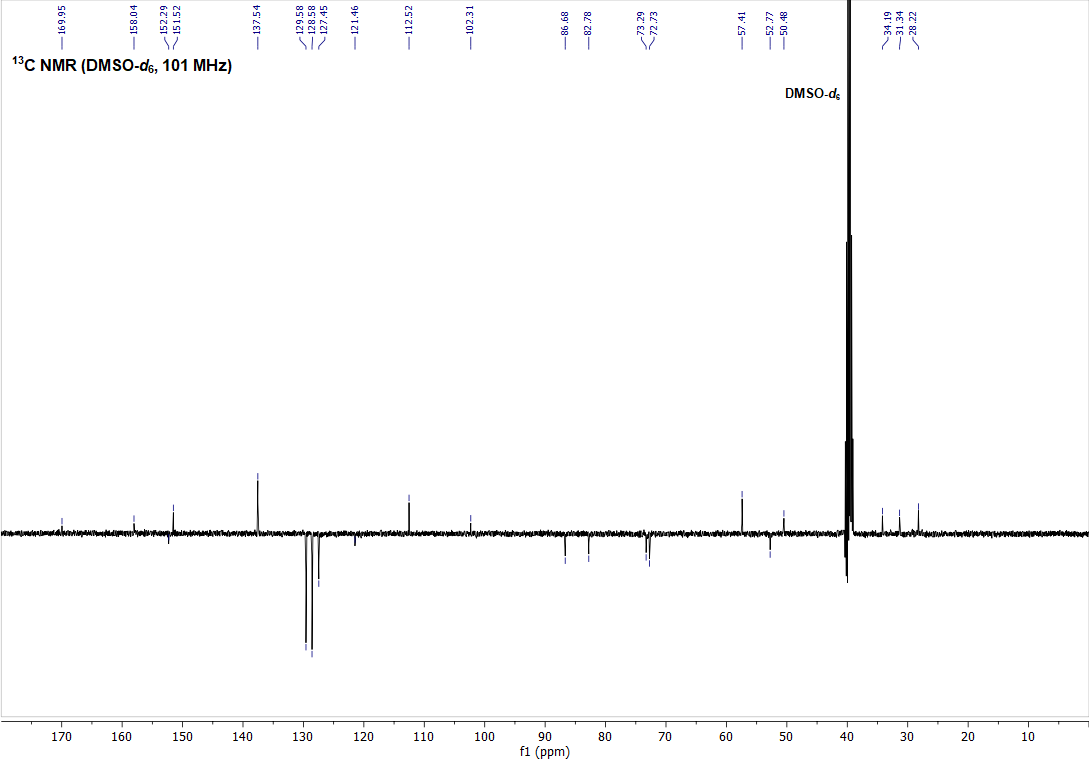

**SI Figure 3** – ^1^H (top) and ^13^C APT (bottom) NMR spectra of compound **STM1078** measured in DMSO-*d*_6_.

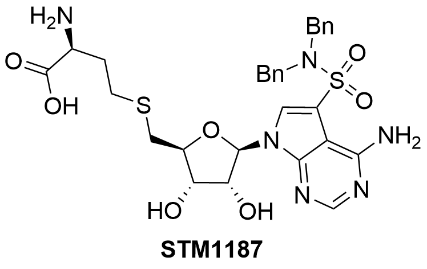

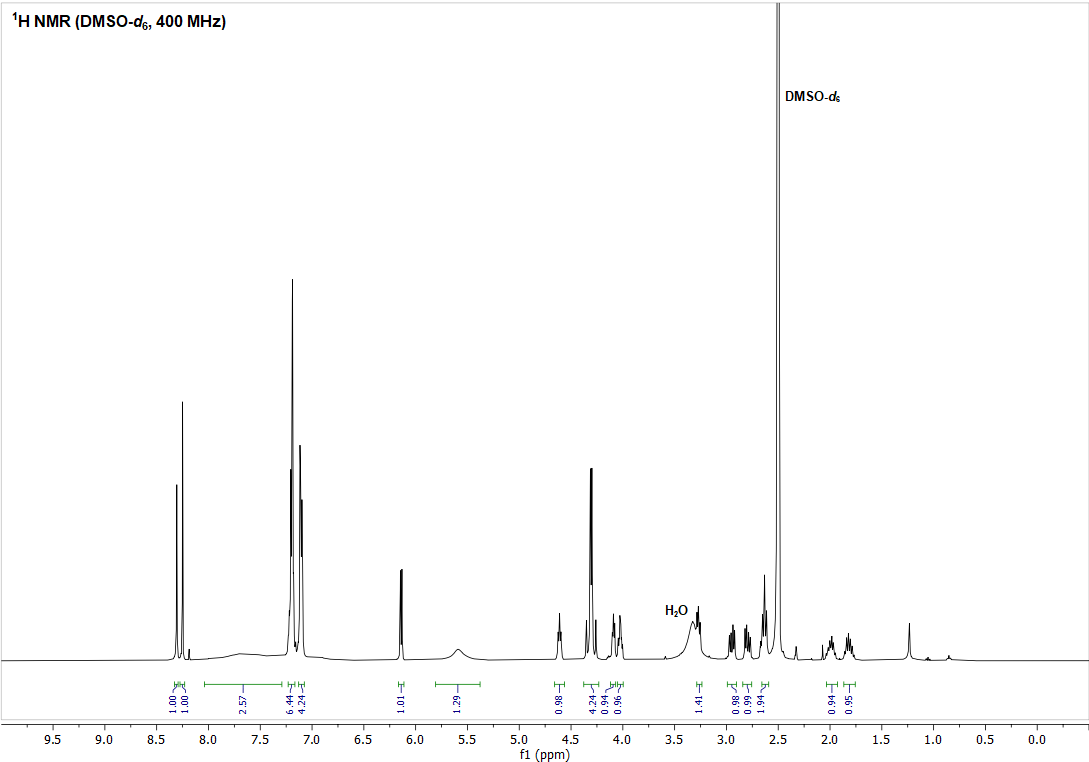

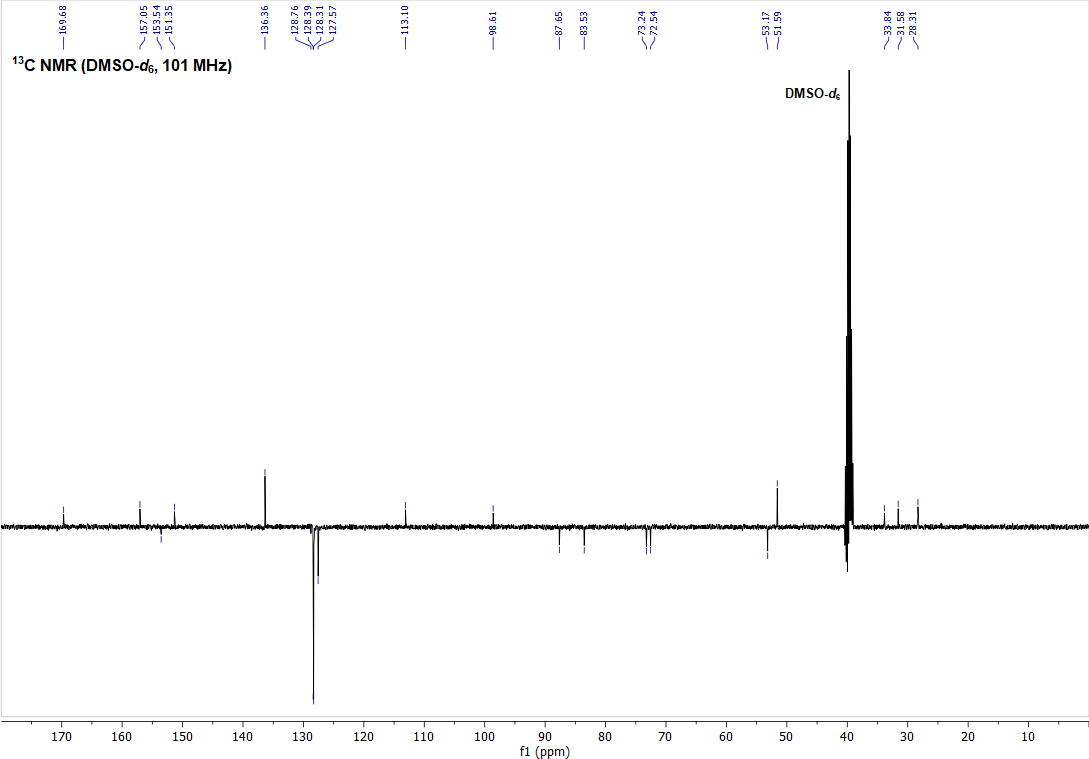

**SI Figure 4** – ^1^H (top) and ^13^C APT (bottom) NMR spectra of compound **STM1187** measured in DMSO-*d*_6_.

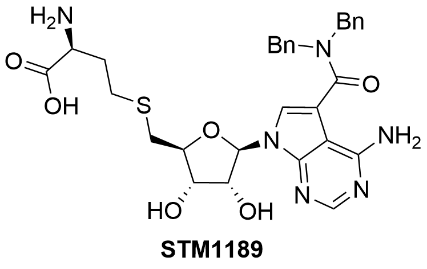

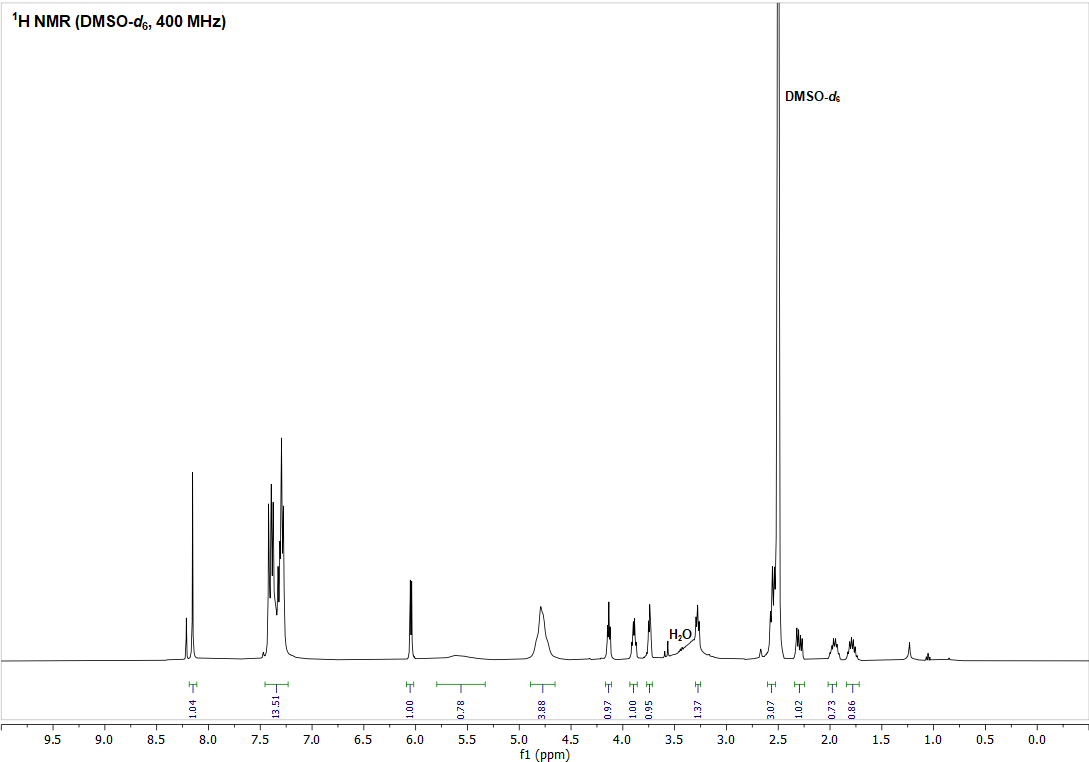

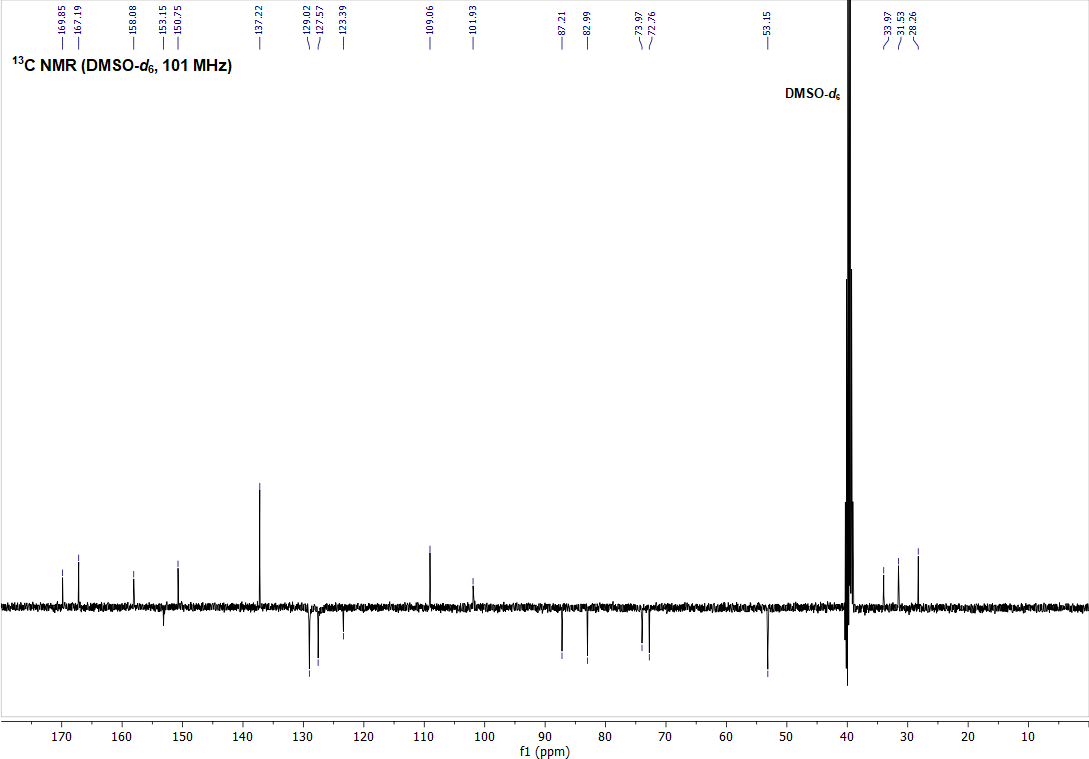

**SI Figure 5** – ^1^H (top) and ^13^C APT (bottom) NMR spectra of compound **STM1189** measured in DMSO-*d*_6_.

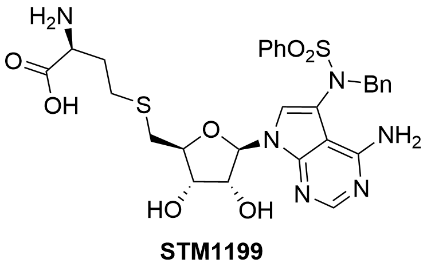

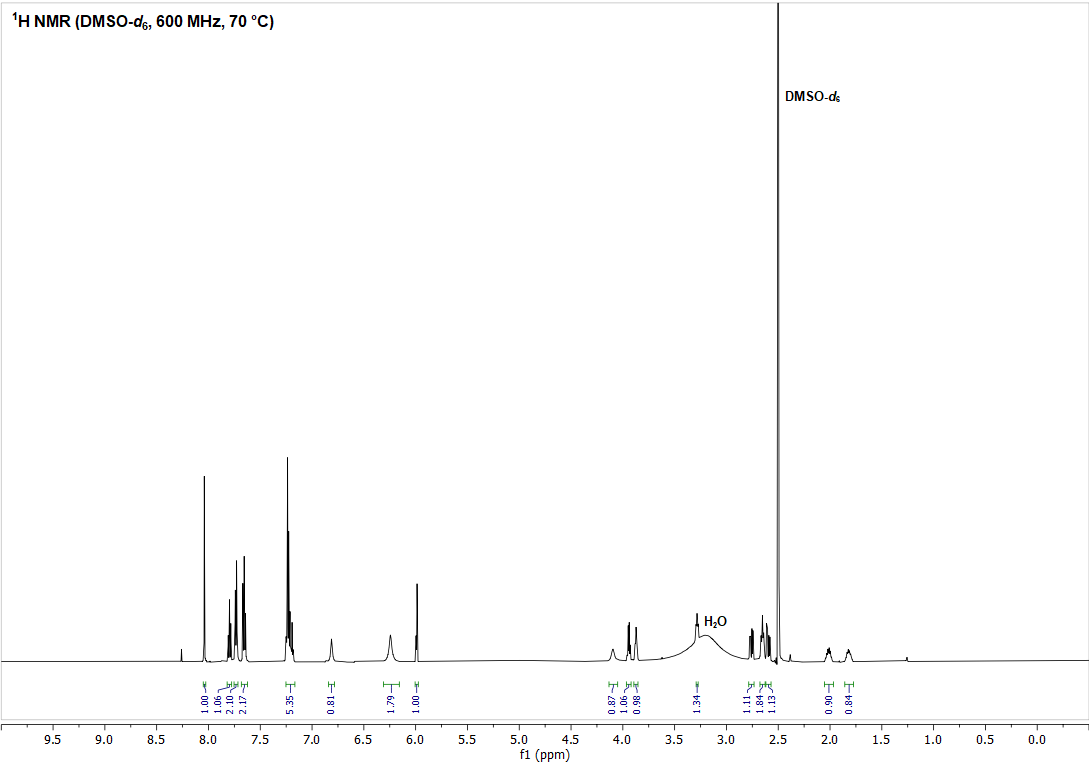

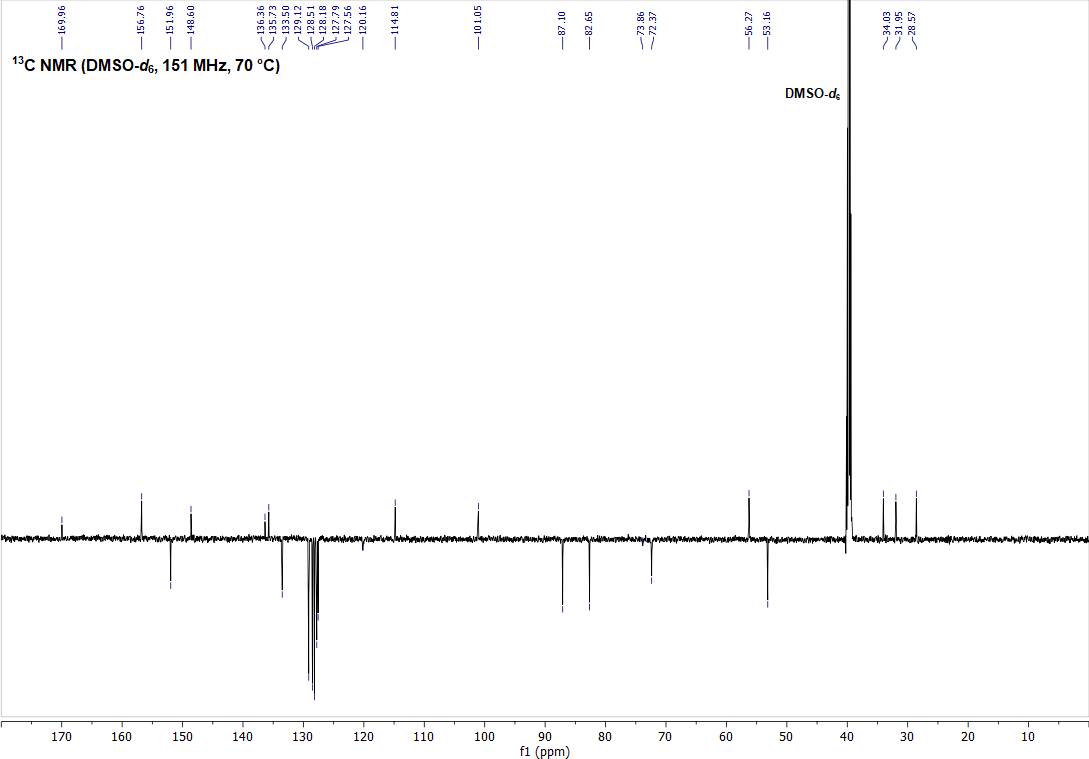

**SI Figure 6** – ^1^H (top) and ^13^C APT (bottom) NMR spectra of compound **STM1199** measured in DMSO-*d*_6_.

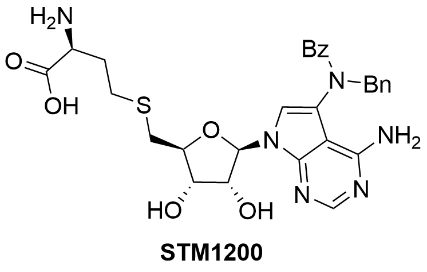

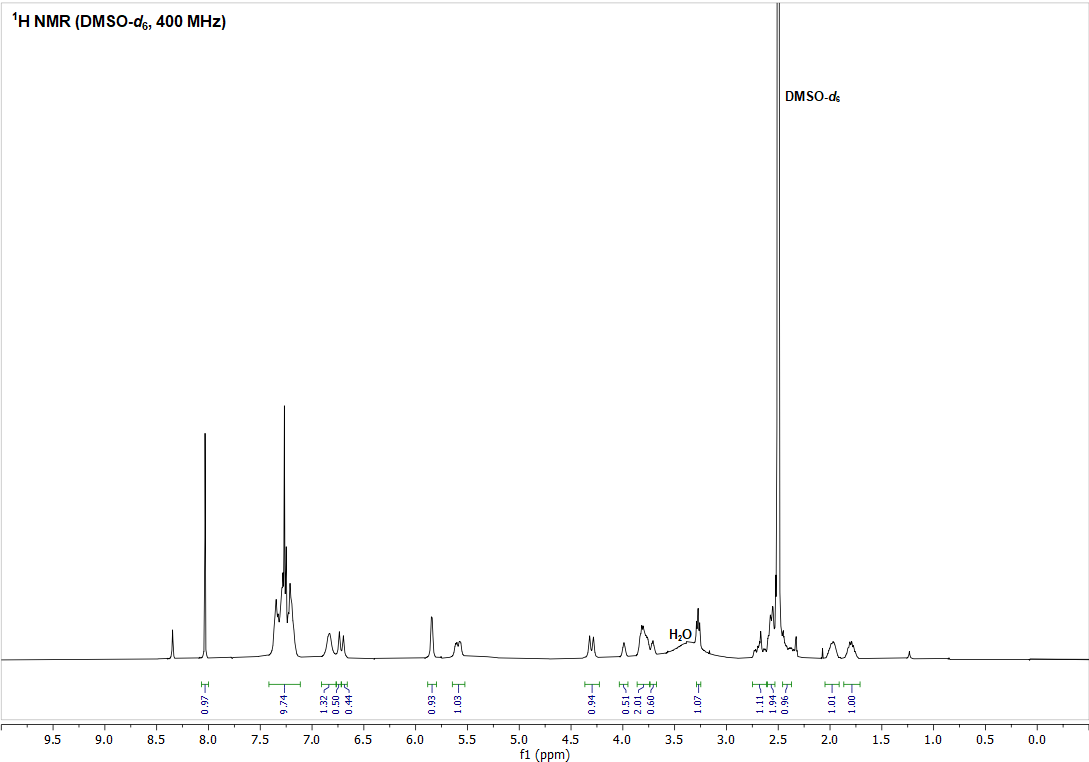

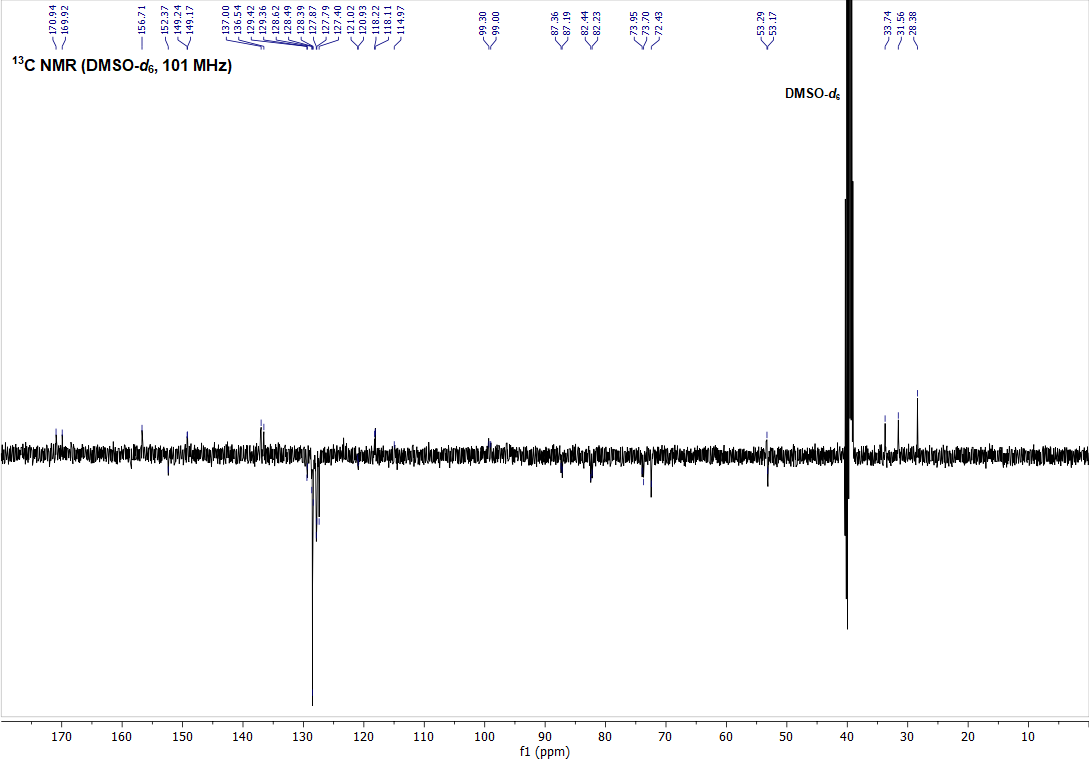

**SI Figure 7** – ^1^H (top) and ^13^C APT (bottom) NMR spectra of compound **STM1200** measured in DMSO-*d*_6_.
